# Glutamate helps unmask the differences in driving forces for phase separation versus clustering of FET family proteins in sub-saturated solutions

**DOI:** 10.1101/2023.08.11.552963

**Authors:** Mrityunjoy Kar, Laura T. Vogel, Gaurav Chauhan, Hannes Ausserwöger, Timothy J. Welsh, Anjana R. Kamath, Tuomas P. J. Knowles, Anthony A. Hyman, Claus A. M. Seidel, Rohit V. Pappu

**Affiliations:** Max Planck Institute of Cell Biology and Genetics, 01307, Dresden, Germany; Department of Molecular Physical Chemistry, Heinrich Heine University, 40225, Düsseldorf, Germany; Department of Biomedical Engineering and Center for Biomolecular Condensates, Washington University in St. Louis, St. Louis, MO 63130, USA; Centre for Misfolding Diseases, Yusuf Hamied Department of Chemistry, University of Cambridge, CB2 1EW, Cambridge, UK

## Abstract

Multivalent proteins undergo coupled segregative and associative phase transitions. Phase separation, a segregative transition, is driven by macromolecular solubility, and this leads to coexisting phases above system-specific saturation concentrations. Percolation is a continuous transition that is driven by multivalent associations among cohesive motifs. Contributions from percolation are highlighted by the formation of heterogeneous distributions of clusters in sub-saturated solutions, as was recently reported for Fused in sarcoma (FUS) and FET family proteins. Here, we show that clustering and phase separation are defined by a separation of length- and energy-scales. This is unmasked when glutamate is the primary solution anion. Glutamate is preferentially excluded from protein sites, and this enhances molecular associations. Differences between glutamate and chloride are manifest at ultra-low protein concentrations. These differences are amplified as concentrations increase, and they saturate as the micron-scale is approached. Therefore, condensate formation in supersaturated solutions and clustering in sub-saturated are governed by distinct energy and length scales. Glutamate, unlike chloride, is the dominant intracellular anion, and the separation of scales, which is masked in chloride, is unmasked in glutamate. Our work highlights how components of cellular milieus and sequence-encoded interactions contribute to amplifying distinct contributions from associative versus segregative phase transitions.

## Introduction

Macromolecular condensation drives the formation of biomolecular condensates that contribute to spatial, temporal, and functional organization of cellular matter ^1, 2^. Condensation combines multiple concentration-dependent processes including associative and segregative phase transitions that include reversible binding, percolation, and phase separation ^3^. The contributions of interactions across difference length and energy scales to the collection of processes that come under the rubric of condensation are coming into focus through systematic investigations ^4, 5, 6, 7, 8, 9, 10^.

The physical chemistry of condensation has been intensively studied using the FET family proteins. These include FUS (Fused in Sarcoma), EWSR1 (Ewing Sarcoma breakpoint region 1/EWS RNA binding protein 1), TAF15 (TATA-Box Binding Protein Associated Factor 15), and at least 25 other proteins ^11^. These proteins feature one or more RNA recognition motifs (RRMs), an intrinsically disordered arginine-rich RNA binding domain (RBD), and a prion-like low complexity domain (PLCD). *In vitro*, in the presence of 150 mM KCl and at a pH of ∼7.2, each FET family protein undergoes phase separation above a sequence-specific and solution-condition dependent saturation concentration (*c*_sat_) ^11^. These saturation concentrations are positively correlated with the endogenous concentrations of FET family proteins ^11, 12^. FET family proteins undergo macrophase separation, with a defined *c*_sat_. However, recent studies showed that FET family proteins also form heterogeneous distributions of clusters in sub-saturated solutions ^7^. These solutions are defined by protein concentrations that are below the system-specific values of *c*_sat_. With increased protein concentration, the average sizes of clusters in sub-saturated solutions grow continuously in size and abundance. The distributions of cluster sizes in sub-saturated solutions are heavy-tailed. Above *c*_sat_, condensate formation via macrophase separation is driven, in part, by cluster-cluster coalescence and the networking of the larger, mesoscopic clusters that form in sub-saturated solutions ^7^.

The sequence determinants of clustering in sub-saturated solutions and *c*_sat_ can be similar ^7^. And yet, clustering in sub-saturated solutions and macrophase separation in supersaturated solutions are also separable processes as evidenced by the fact that of solutes such as ATP and 1,6-hexanediol dissolve condensates, but do not have an effect on clusters in sub-saturated solutions ^7^. Further while certain mutations impact clustering, they have a minimal impact on phase separation. These observations suggest that clustering and phase separation are distinct processes governed by separable length and energy scales. Does this separation prevail in cellular milieus? If yes, it would imply that clusters that form in sub-saturated solutions are likely to prevail at endogenous or lower than endogenous levels of expression. To answer this question, we investigated the effects of glutamate on the driving forces for macrophase separation versus cluster distributions in sub-saturated solutions.

In cellular milieus, the solvent is a complex mixture of ions, metabolites, and osmolytes ^13, 14^. The high concentrations of potassium (∼150 mM) are balanced by anions that include glutamate, glutathione, and organic phosphates ^14^. Importantly, the relevant anion inside cells is glutamate, whereas the intracellular concentrations of chloride are very low by comparison ^14, 15, 16^. The concentrations of glutamate vary across cell types, but it is usually in the 10 - 100 mM range ^13, 14^. There are important physicochemical differences between chloride and glutamate. For chloride, the charge of the anion is localized to the Cl atom (**Fig. 1a**), and its pK_a_ is ∼ −4, making it a very strong acid or even a weak base ^17^. In contrast, the pK_a_ of the free carboxylate anion of glutamate is ∼4.8, making it a weak acid (**Fig. 1b**). The charge on the carboxylate moiety is delocalized across the two *sp*^2^ hybridized oxygen atoms. Further, the ionic potential value of Cl^-^ is higher than the carboxylate anion. As a result, chloride polarizes cations more strongly than the carboxylate anions. These details are likely to have a direct impact on the effective strengths of protein-protein and protein-nucleic acid interactions ^18, 19, 20, 21^. Recent studies have shown that protein-protein interactions of bacterial single-stranded DNA binding proteins (SSBs) are stronger in glutamate when compared to chloride ^22, 23^. Further, the cooperativity of binding of bacterial SSBs to their nucleic acid substrates, as influenced by the intrinsically disordered C-terminal tails, is readily unmasked in the presence of glutamate as opposed to chloride ^24^.

**Figure 1:**
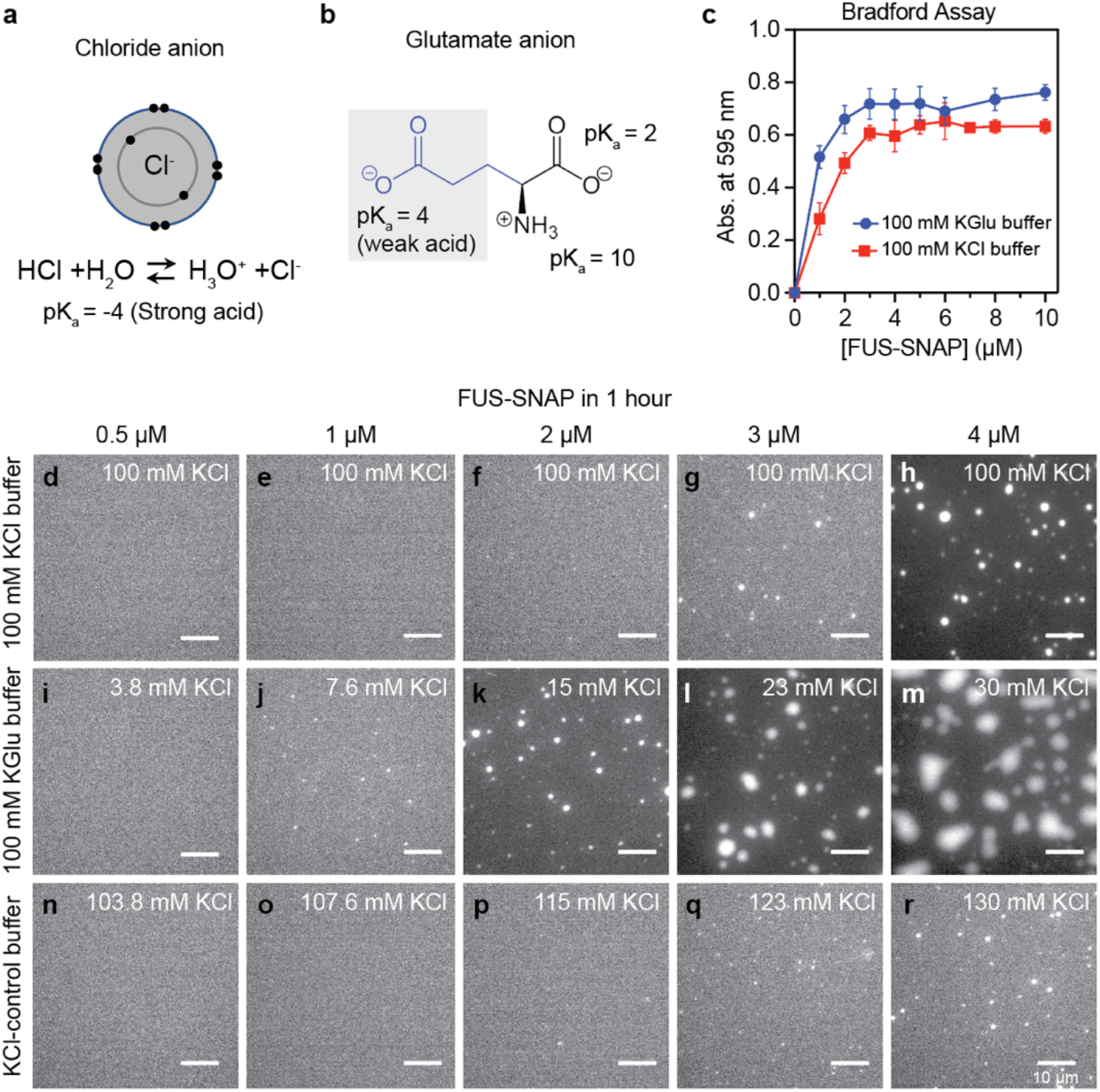
The *c*sat of FUS-SNAP is similar in KCl versus KGlu buffers. The timescales for forming micron-scale condensates shows differences influenced by clustering at concentrations below *c*sat. (a) Schematic of chloride anion showing its electron distribution in 2*s* and 2*p* orbitals as well as its pKa. (b) Schematic of glutamate. (c) Sample data for absorbance-based spin-down assays. Data are shown here for FUS-SNAP in 20 mM Tris.HCl pH 7.4, with 100 mM KCl and 20 mM Tris.Glu pH 7.4 and 100 mM Kglu at ≈ 25°C. Panels (d)-(h) show bright field microscopy images collected at the 1-hour time point for solutions containing different concentrations of FUS-SNAP in 20 mM TRIS.HCl, pH 7.4, with a final KCl concentration of 100 mM. Panels (i)-(m) show microscopy images collected at the 1-hour time point for solutions containing different concentrations of FUS-SNAP in 20 mM Tris.Glu, pH 7.4, with 100 mM KGlu, and panels (n)-I show microscopy images collected at the 1-hour time point for solutions containing different concentrations of FUS-SNAP in 20 mM Tris.HCl, pH 7.4, with 100 mM KCl. In both KGlu and KCl-control buffer, the residual KCl (< 30 mM) from FUS-SNAP stock was added. The total concentration of KCl is marked on the panels (i-r). For imaging purposes, 5% of the total mixture in each sample is made up of Alexa 488 labeled FUS-SNAP. The scale bar in each panel corresponds to 10 µm.

Here, we investigate the effects of glutamate on the clusters formed by FET family proteins in sub-saturated solutions. Changing the dominant anion from chloride to glutamate has minimal effects on the driving forces for phase separation. However, the presence of glutamate significantly enhances the extent of clustering in sub-saturated solutions. These results emphasize the separation of energy scales that contribute to associations in sub-saturated solutions versus the driving forces for phase separation and highlight the fact that this separation is amplified in conditions that are closer mimics of cellular milieus.

## Results

### Enabling assessments of the effects of glutamate versus chloride

The FET family proteins cannot be purified or stored in buffers containing potassium glutamate (KGlu) because the proteins readily precipitate under these conditions. Therefore, the proteins were purified and stored in a buffer with a high concentration of potassium chloride (KCl) (50 mM Tris-HCl pH 7.4, 500 mM KCl, 5% Glycerol, and 1 mM DTT) ^7^. To prepare solutions at different protein concentrations in 100 mM KCl, the stock protein solutions were diluted and adjusted with KCl buffer. To prepare protein solutions in 100 mM KGlu, the protein stock solution was diluted with 100 mM KGlu buffer. Accordingly, these solutions contain an additional ∼1 mM to ∼30 mM KCl from the stock protein solution. To assess the effects of the residual KCl from the stock protein solutions in KGlu buffers, we performed control measurements in conditions referred to as “KCl-control buffer.” This includes the corresponding amount of residual KCl from the stock protein solution and 100 mM KCl.

### Saturation concentrations of FUS change minimally in glutamate versus chloride buffers

Using a spin-down-based absorbance assay, we quantified saturation concentrations (*c*_sat_) for SNAP (**S**y**n**aptosmal-**A**ssociated-**P**rotein)-tagged FUS, designated as FUS-SNAP. The values of *c*_sat_ were measured in 100 mM KGlu and 100 mM KCl. The spin-down assay yields an estimate of 2 µM and 3 µM for the *c*_sat_ of FUS-SNAP in 100 mM KGlu versus 100 mM KCl, respectively (**Fig. 1c**).

Next, we collected microscopy images ∼1 hour and ∼4 hours after sample preparation. Results at the 1-hour time point are shown in **Fig. 1d-1r** for different concentrations of FUS-SNAP in 100 mM KCl, 100 mM KGlu, and the KCl-control buffer. In 100 mM KCl or KCl-control buffer, the formation of micron-scale condensates is detectable only at or above ∼3 µM. However, in 100 mM KGlu, we observed small puncta in the concentration range of 1-2 µM. These puncta do not grow significantly over the period of four hours (**Extended Data Fig. 1**). Increasing the protein concentration from 1 µM to 4 µM leads to the growth of micron-scale puncta over the four-hour window. These data suggest that glutamate has a minimal effect on the location of the phase boundary when compared to chloride. However, it appears to influence the extent of clustering in sub-saturated solutions, which in turn impacts the differential kinetics for phase separation in supersaturated solutions (see data in **Fig. 1d-1r** and **Extended Data Fig. 1**).

### Independent assessments of *c*_sat_ estimates

We performed dynamic light scattering (DLS) in different solutions containing different amounts of FUS-SNAP to obtain an independent assessment of the estimates of *c*_sat_ for macrophase separation (**Extended Data Fig. 2**). DLS helps probe the presence of slow modes in the temporal evolution of autocorrelation functions, the onset of which is a feature of supersaturated solutions ^7^. In 100 mM KCl, the autocorrelation functions reach a steady state below *c*_sat_ (< 3 µM) and do not change significantly over 26 minutes (Fig. 1d**, 1e**). Above *c*_sat_ (**Extended Data Fig. 2c**), the autocorrelation functions show the presence of slow modes, which are signatures of coalescence and networking of clusters that drive macrophase separation ^7^. In KGlu, the slow modes appear at protein concentrations of 2 µM (**Extended Data Fig. 2e**), whereas in KCl they appear at 3 µM ^7^. These results provide independent confirmation of the estimates of *c*_sat_ and emphasize the small differences in driving forces for macrophase separation of FUS-SNAP in 100 mM KCl versus 100 mm KGlu.

### Clustering of FET proteins in sub-saturated solutions is enhanced in glutamate when compared to chloride

The driving forces for macrophase separation, which refers to the formation of micron-scale condensates, is minimally affected by glutamate when compared to chloride. However, we know, from previous studies, that FET family proteins also form heterogeneous distributions of clusters in sub-saturated solutions ^7^. Theory, simulation, and extant data suggest that clustering in sub-saturated solutions and macrophase separation can be governed by distinct energy scales ^7, 25^. The recent work of Lan et al., also affirms these findings ^8^, albeit for different systems. Given the likely differences between preferential interactions of chloride versus glutamate on the molecular scale ^18, 20, 21^, we asked if the two anions differently influence clustering in sub-saturated solutions.

We used nanoparticle tracking analysis (NTA) ^26^ to quantify the concentration dependence of forming mesoscale clusters in sub-saturated solutions (**Movies S1** and **S2**). Mesoscale clusters are defined as species that are hundreds of nanometers in diameter and are large enough to be detected by light scattering ^7^. We collected NTA data for FUS-SNAP and untagged FUS in 100 mM KGlu and 100 mM KCl. At equivalent protein concentrations, the abundance of mesoscale clusters is ∼4-fold higher in 100 mM KGlu when compared to 100 mM KCl (Fig. 2a**-2c** and. **Extended Data Fig. 3**). This translates to an advantage of ∼0.5 – 0.75 kcal/mol in the free energy of transfer of FUS molecules from the dispersed phase into mesoscale clusters in glutamate versus chloride.

**Figure 2:**
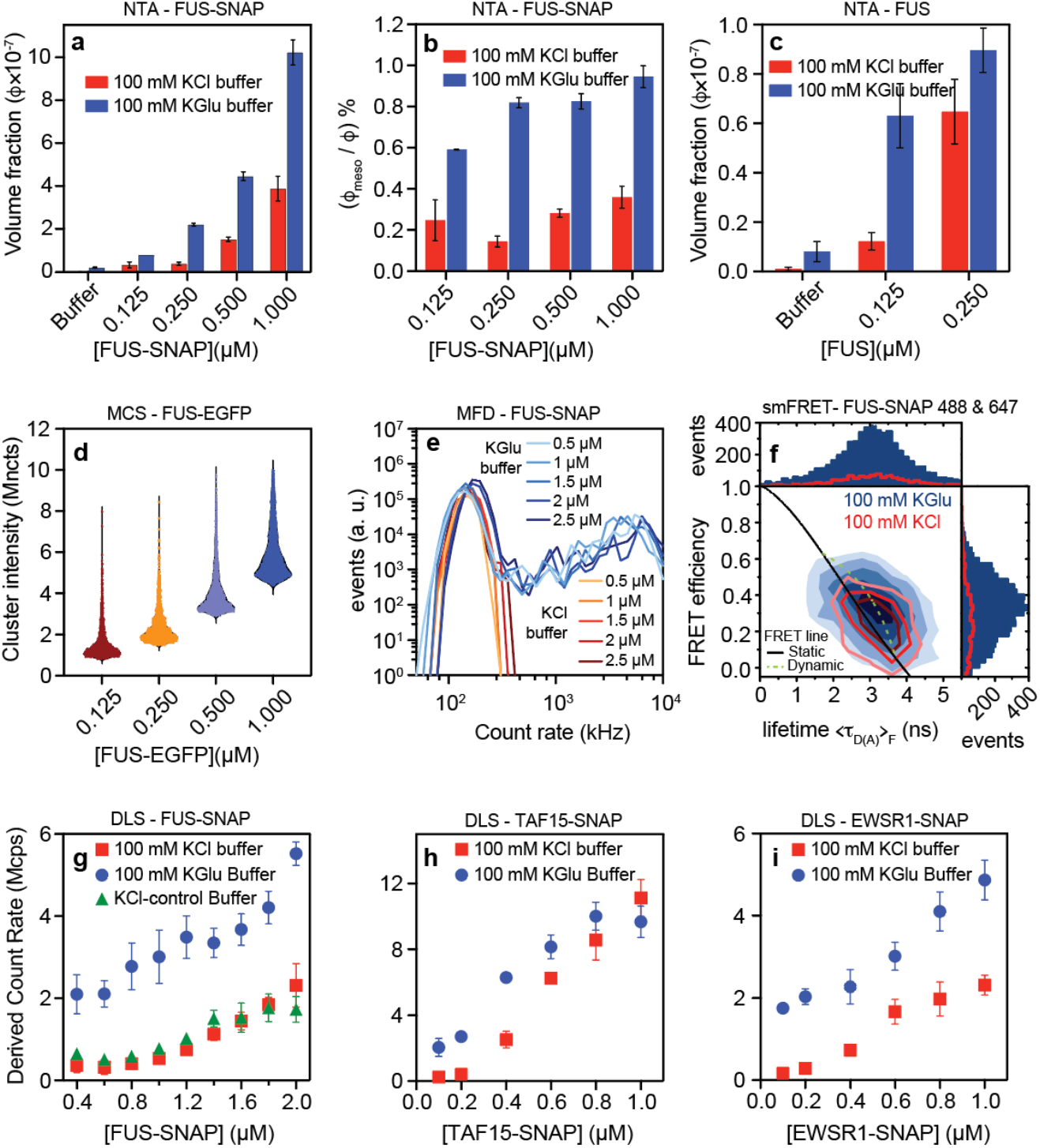
The abundance of mesoscale clusters in FUS and FUS family proteins increases in KGlu buffers compared to KCl buffers. (a) NTA data shows the volume fraction of mesoscale clusters for FUS-SNAP at different sub-saturation concentrations in 100 mM KCl and KGlu buffers. (b) Relative abundance of mesoscale clusters for FUS-SNAP at different sub-saturation concentrations in 100 mM KCl and KGlu buffers. (c) NTA data shows the volume fraction of mesoscale clusters for FUS at different sub-saturation concentrations in 100 mM KCl and KGlu buffers. (d) Single-molecule analysis by microfluidic confocal spectroscopy (MCS) shows that FUS-EGFP cluster formation increases with increasing concentration from 125 nM to 1000 nM in KGlu buffer. (e) The intensity distributions from the multiparameter fluorescence detection (MFD) experiment in the presence of 15 nM Nile red titrated with various concentrations of FUS-SNAP shows that the amount and size of cluster increase in 100 mM KGlu visibly by tailing towards higher count rates up to 10^4^ kHz per burst compared to 100 mM KCl buffer. (f) Single-molecule FRET measurements with 200 pM of FUS-SNAP-AF488 as donor and FUS-SNAP-AF647 as acceptor in KGlu buffer (blue) compared to KCl buffer (red) show that the number of FRET events are significantly higher for KGlu (1D-histograms). The FRET populations are broadened beyond shot noise due to intermolecular dynamics (dynamic FRET line, dashed light green), resulting in a deviation from the static FRET line (solid black). DLS data shows the derived count rate of FUS-SNAP (d), TAF15-SNAP (e), and EWSR1-SNAP (f) in different buffers.

Next, we used microfluidic confocal spectroscopy (MCS) to investigate the totality of cluster sizes that form in sub-saturated solutions in the presence of glutamate. MCS is a brightness-based method that combines microfluidic mixing and flow with confocal detection ^27, 28, 29^. As with previous studies, which were performed in the presence of 100 mM KCl ^7^, we observed a monotonic increase in the brightness per molecule for FUS-EGFP in 100 mM KGlu where EGFP refers to enhanced green fluorescent protein. This points to the formation of clusters in sub-saturated solutions whose average sizes increase with increasing concentrations (Fig. 2d).

Building on the MCS measurements, we used multiparameter fluorescence detection (MFD) as a complementary method that affords access to the distributions of clusters that form in the presence of glutamate versus chloride. In these experiments, we used increasing concentrations of FUS-SNAP stained with a fixed concentration (15 nM) of Nile Red in the presence of either 100 mM KGlu or 100 mM KCl. The fluorescence intensity distributions, which were extracted using analysis of fluorescence bursts, show clear and pronounced differences between glutamate versus chloride (Fig. 2e). Additionally, in glutamate, we observed a continuous tailing towards count rates of up to 10^4^ kHz. In contrast, we did not observe significant populations beyond 400 kHz for FUS-SNAP in KCl. These results suggest that clustering in sub-saturated solutions is significantly enhanced in the presence of glutamate versus chloride.

Fluorescence intensity distribution analysis (FIDA) provides insights regarding the growth in brightness, and this is a useful proxy for the sizes of protein clusters. Inferences regarding the sizes of protein clusters can be drawn by analysis of donors and acceptors separately (**Extended Data Fig. 3**). The elongated shape of the contours in two-dimensional histograms (Fig. 2f) suggests the formation of dynamic clusters of FUS molecules that rearrange on the millisecond timescale. This is supported by clear deviations from the static FRET line. Assuming a simple structural model consisting of two states, the datasets from both KCl and KGlu buffers can be fitted with the same dynamic FRET line. Further, measurements of fluorescence correlation spectroscopy (FCS) on the single-molecule level show that the autocorrelation of donor-labelled FUS-SNAP is consistent with increased translational diffusion time t_d2_ and strong fluctuation in the weighted residuals at correlation times longer than t_d2_ in the KGlu buffer (**Extended Data Fig. 3f**). This points to the presence of higher-order complexes even at 400 pM concentrations of FUS-SNAP. These single-molecule data show clear evidence for increased heterogeneity and larger cluster sizes for FUS-SNAP in glutamate versus chloride.

Next, we used DLS to examine the relative abundance of cluster populations in different buffers by measuring the derived count rate, which is the theoretical count rate one would obtain at 100% laser power with zero attenuation. The higher derived count rate indicates higher abundance and larger particles. Consistent with NTA data, the derived count rate for FUS-SNAP is up to 3-fold higher in 100 mM KGlu when compared to 100 mM KCl (Fig. 2g). Untagged FUS shows similar behaviors (**Extended Data Fig. 3g**).

To assess the transferability of our findings to other members of the FET family, we performed DLS measurements and quantified derived count rates for FUS-EGFP (**Extended Data Fig. 3h**), TAF15-SNAP (Fig. 2h) and EWSR1-SNAP (Fig. 2i). In all systems, we observe a similar trend, where the derived count rate is higher in 100 mM KGlu when compared to 100 mM KCl at each of the protein concentrations studied. The protein concentration of TAF-SNAP in its stock solution is low (∼12 µM). Accordingly, with increasing protein concentrations, the method of sample preparation leads to higher residual KCl in the solution. Therefore, at 100 mM KGlu buffer, the residual KCl suppresses some of the clustering, resulting in a lower difference in the derived count rate than in 100 mM KCl. Similar effects were also observed for untagged FUS (**Extended Data Fig. 3g**). Overall, our measurements suggest that the formation of clusters in sub-saturated solutions is enhanced in 100 mM KGlu compared to 100 mM KCl. This suggests that even though *c*_sat_ changes minimally when the solution anion is glutamate versus chloride, the molecular-scale associations in sub-saturated solutions are stronger in glutamate when compared to chloride.

### Differential effects of glutamate versus chloride are evident in their influence on mutations within FUS

We investigated the effects of mutations within the FUS PLCD on clustering in sub-saturated solutions in the presence of 100 mM KGlu versus 100 mM KCl buffer. Tyr-to-Ser mutations increase *c*_sat_ by up to two orders of magnitude ^11^. Mesoscale clusters, as measured by the derived count rates, are undetectable in either 100 mM KCl or 100 mM KGlu. However, while substitution of 27 Tyr residues to Ser in the PLCD makes mesoscale clustering undetectable via DLS, some level of clustering is evident when 18 or 10 Tyr residues in the PLCD are substituted to Ser in the presence of 100 mM KGlu (Fig. 3a). These results implicate the Tyr residues in PLCDs as being important for clustering and phase separation. These interactions appear to be strengthened in glutamate. Note that the DLS measurements for Tyr-to-Ser mutations in the PLCD of FUS were performed at protein concentrations of 10 µM, which is almost two orders of magnitude higher than the concentrations used for wild-type FUS. Substitution of 24 Arg residues within the RBD to Gly also increases *c*_sat_ by up to two orders of magnitude compared to wild-type FUS ^11^. These substitutions abrogate clustering in sub-saturated solutions in both chloride and glutamate (Fig. 3b). Taken together with the results from substitutions of 27 Tyr to Ser in the PLCD, the substitutions of Arg to Gly in the RBD emphasize the importance of networks of Tyr-Arg interactions as drivers of phase separation and clustering in sub-saturated solutions.

**Figure 3:**
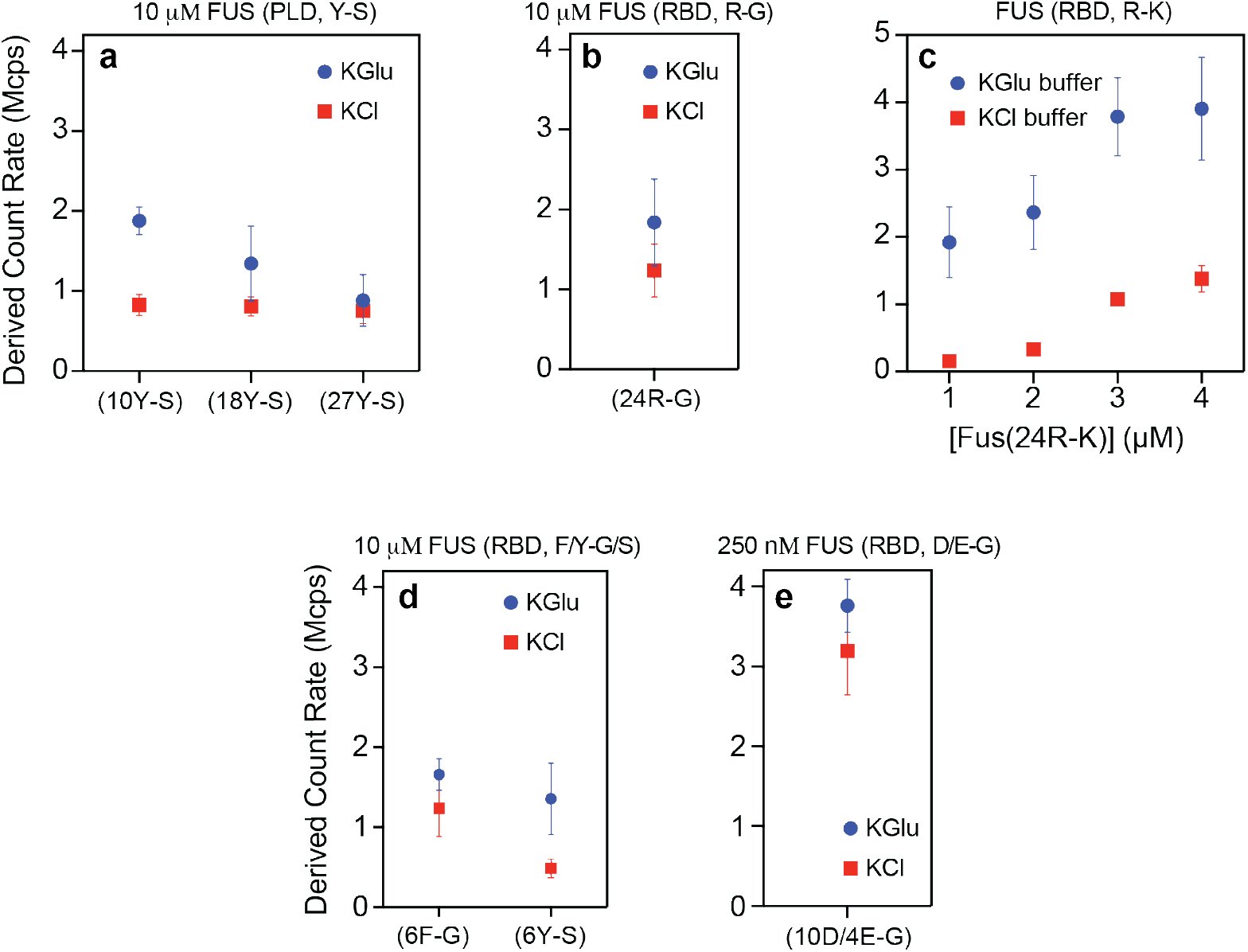
Mutations modulate the extent of clustering in sub-saturated solutions and this can be influenced by glutamate. DLS data show the derived count rate of FUS(Y-S) (a), FUS(R-G) (b), FUS(R-K) (c), FUS(F/Y-G/S) (d), and FUS(10D/4E-G) (e) in different buffers.

Substituting 24 Arg residues in the RBD to Lys increases *c*_sat_ by an order of magnitude compared to wild-type FUS ^11^. Interestingly, while these substitutions weaken clustering in the presence of 100 mM KCl (Fig. 3c), the extent of clustering we observe in the presence of 100 mM KGlu is similar to that of wild-type FUS (compare Fig. 3c to **Extended Data Fig. 3g**). As shown in recent single molecule studies, there is a uniform weakening of cation-π interactions in chloride salts ^30^. In contrast, in glutamate the differences between wild-type FUS and the 24R-K variant seem to be length-scale dependent. Specifically, while clustering, which is mainly impacted by the network of cation-π interactions, is preserved upon substituting Arg to Lys, phase separation, which should be governed mainly by solubility, is weakened considerably by Arg to Ly substitutions. This can be rationalized if the strengths of cation-π interactions are minimally affected by glutamate when compared to chloride. However, the increase in *c*_sat_ points to differences in solubility driven by substitutions of Arg to Lys. Indeed, it is noteworthy that the free energies of solvation of Arg and Lys are fundamentally different from one another ^31^. From this hydration perspective, Arg has more of a hydrophobic character than Lys ^31^. As a result, the driving forces for phase separation, which are governed mainly by solubility, are affected by the substitution of Arg to Lys are affected significantly.

The importance of networks of cation–π interactions on clustering in sub-saturated solutions is also underscored by the effects of substitutions of Tyr or Phe residues within the RBD to Ser or Gly, respectively. In both chloride and glutamate, these substitutions weaken the extent of clustering in sub-saturated solutions (Fig. 3d). Finally, substitutions of 10 Asp and 4 Glu residues within the RBD significantly enhance clustering compared to wild-type FUS, and the effects of these substitutions are similar in glutamate and chloride (Fig. 3e). Note that while clustering is weakened and its detection requires the use of 10 µM protein for the 6F-G and 6Y-S variants (Fig. 3d), clustering is readily apparent at 250 nM of the 10D/4E-G variant (Fig. 3e).

### Clusters form reversibly in sub-saturated solutions

The average sizes of clusters in sub-saturated solutions from DLS experiments show a decrease upon dilution and an increase with increased concentration (Fig. 4a**, 4b**). This is true in glutamate and chloride buffers ^7^. Similarly, in MFD experiments (Fig. 4c), when 3 μM of Nile Red stained FUS-SNAP was diluted to 1 μM, the number of bursts decreased. Increasing the protein concentration to 2.75 μM also increased the bursting, and upon dilution to 1.65 μM, the number of bursts decreased. For each protein concentration, the increases in fluorescence amplitude and broadening of the fluorescence distributions become less significant at later dilution steps indicating the reversibility of FUS complexes on the molecular scales while mesoscale clusters remain in solution. In smFRET studies, the introduction of 40 nM of unlabeled FUS-SNAP causes a shift in the FRET signal to very low FRET between FUS-SNAP-AF488 and FUS-SNAP-AF647 (**Extended Data Fig. 4a**). This suggests that cluster formation is reversible and that the molecules within clusters are labile in 100 mM KGlu as well as in chloride ^7^.

**Fig. 4:**
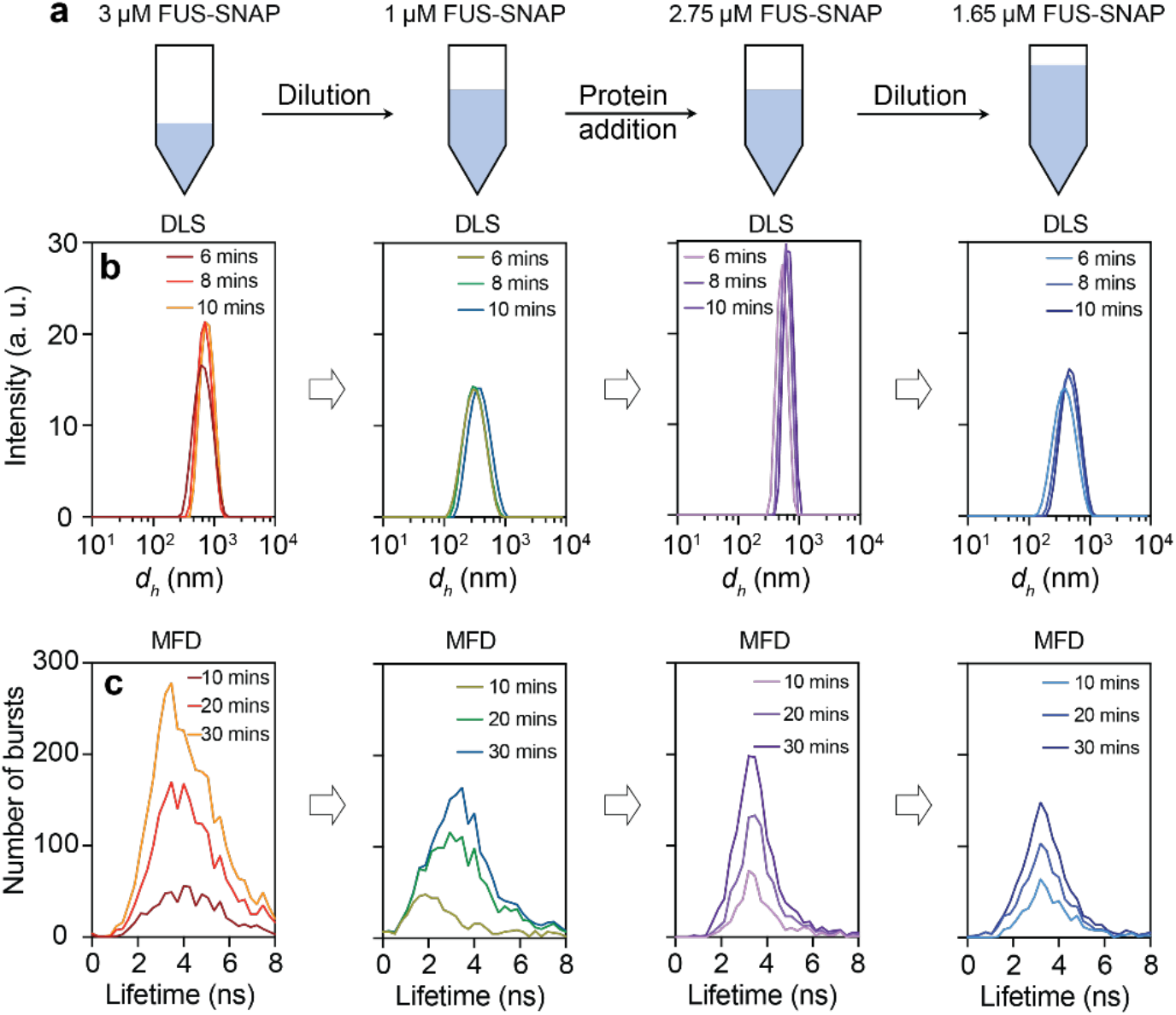
FUS-SNAP clusters form reversibly in the 100 mM KGlu buffer. (a) A schematic representation of the reversibility experiments of FUS-SNAP clusters in KGlu buffer. (b) DLS data were collected at different time points for FUS-SNAP, showing the changes in intensity vs. size distribution profiles at 3 µM, 1 µM, 2.75 µM, and 1.65 µM. (c) The confocal point measurements with multiparameter fluorescence detection (MFD) display the number of burst vs. fluorescence lifetime of Nile Red at the same concentrations shown in (b), revealing repeatedly shifts to lower lifetimes upon dilution.

### Clusters create unique local environments

Next, we used environmentally-sensitive dyes, specifically Nile red and bis-ANS, to probe the physicochemical properties of clusters that form in different buffers. Nile Red shows increased quantum yields in nonpolar environments ^32^. We measured the fluorescence lifetime distributions of Nile red using MFD in various concentrations of FUS-SNAP in 100 mM KGlu and 100 mM KCl. Nile Red exhibits a spectrum of lifetimes ranging from 0.6 ns to 4.66 ns, and the lifetimes increase with increased hydrophobicity ^33^. In 100 mM KCl, at 0.5 μM FUS-SNAP, the peak in the fluorescence lifetime distribution occurs at 2 ns. With increasing concentration of FUS-SNAP, the lifetime distribution in 100 mM KCl becomes bimodal, showing peaks at 2 ns and 4 ns. In the presence of 100 mM KGlu, the Nile Red lifetime distributions show one distinct peak with an average lifetime of 4 ns which is reached at FUS-SNAP concentrations as low as 0.5 µM. The inference is that there is an increase in the number and size of clusters in KGlu when compared to KCl (Fig. 5a).

**Fig. 5:**
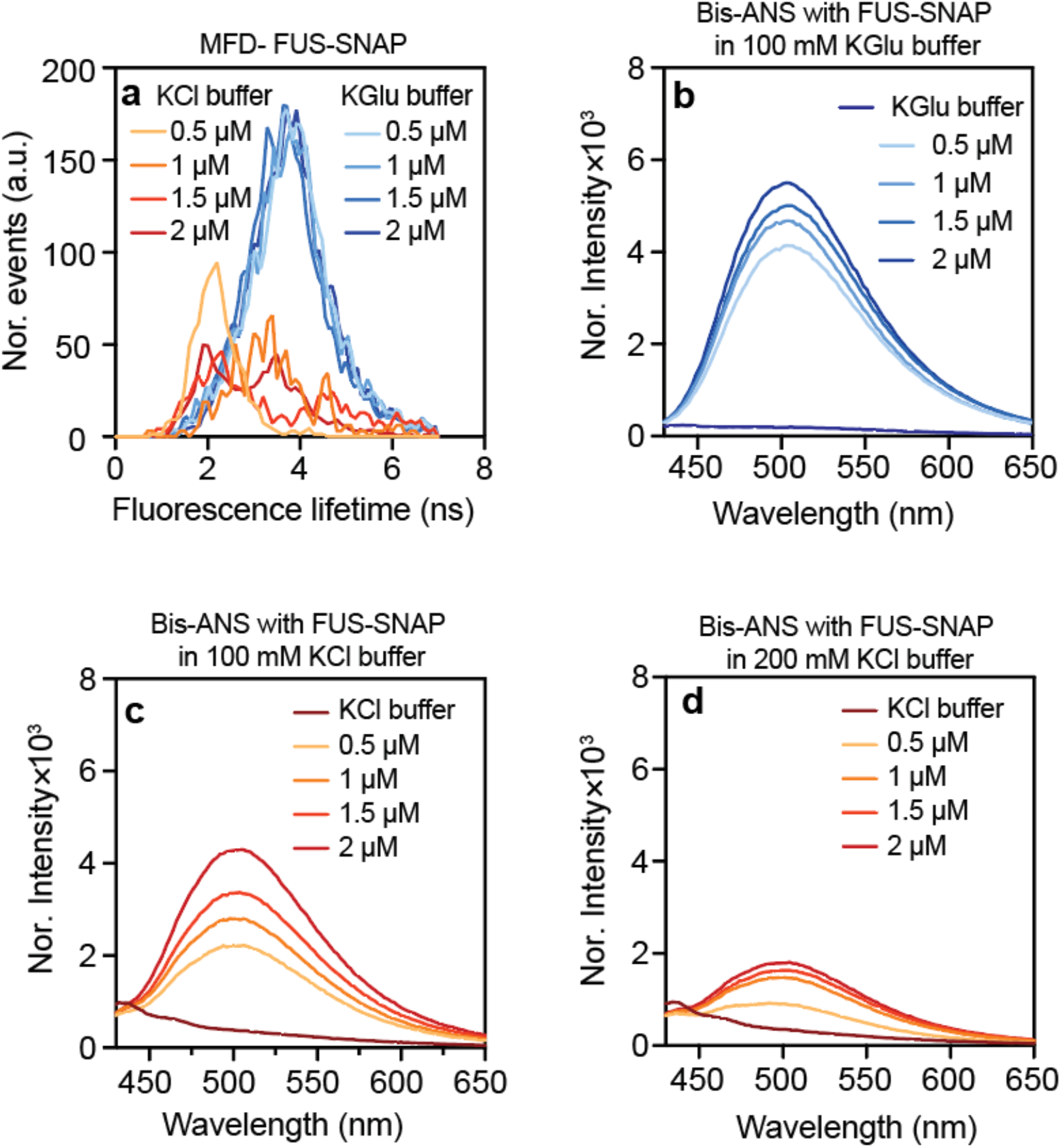
FUS-SNAP clusters create a distinct microenvironment compared to proteins in solution. (a) The fluorescence lifetimes of Nile Red with various concentrations of FUS-SNAP equilibrated for 30 minutes in KGlu and KCl buffer. (b)-(d) The different concentrations of FUS-SNAP solutions and buffers are mixed with 2 µM bis-ANS in 100 mM KGlu buffer (b), 100 mM KCl buffer (c), and 200 mM KCl buffer (d). For bis-ANS studies (b)-(d), the mixture solutions were excited using a 355 nm laser, and the emission spectra were measured from 425 nm to 650 nm.

We also used bis-ANS to probe the local environments within clusters that form in the presence of 100 mM KGlu versus 100 mM KCl (Fig. 5b and 5c). In both cases, in the presence of a constant 2 μM bis-ANS concentration, the fluorescence intensity increases with increasing protein concentration. The increase in intensity is higher in the presence of KGlu compared to KCl. As a control, when we increased the KCl concentration to 200 mM, it caused a decrease in the fluorescence intensity of bis-ANS with FUS-SNAP compared to 100 mM KCl buffer. This suggests that KCl inhibits the clustering of FUS-SNAP. To assess the hydrophobicity of the clusters, the fluorescence intensity of the same concentration of bis-ANS was measured in solvents, methanol, and ethanol (**Extended data** Fig. 4b). According to the data, the intensity of bis-ANS in the presence of FUS-SNAP clusters in KGlu buffer is comparable to that of bis-ANS in methanol. The dielectric constant of methanol is smaller by a factor of 0.4 when compared to that of water at 25°C ^34^. Overall, our findings suggest that clustering in sub-saturated solutions is enhanced in KGlu when compared to KCl. Stronger molecular associations increase the extent of clustering, the sizes of clusters, and the larger clusters are more hydrophobic when compared to the surrounding solvent.

### Glutamate is preferentially excluded from amides

To uncover the molecular basis for the differences in cluster formation in KGlu versus KCl, we studied the interactions of the two salts with different amino acids using molecular dynamics (MD) simulations. These simulations use capped amino acids and explicit representations of solvent molecules and solution ions. The simulation results were used to compute preferential interaction coefficients for the two salts for different amino acids (Glycine, Lysine, Arginine, Serine, Aspartic acid, and Glutamine).

Preferential interaction coefficients quantify the excess numbers of cosolutes in the vicinity of the peptide when compared to the bulk solution. Using local-bulk partitioning ^35^ (Fig. 6a), the coefficients were calculated as: 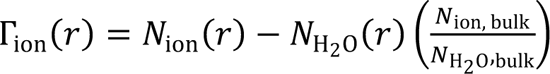. Here, *N*_ion_(γ) and *N*_H_2_O_(γ) quantify the numbers of ions and water molecules at a distance *r* from the center-of-mass of the peptide, whereas *N*_ion, bulk_ and *N*_H2O_,_bulk_ quantify the numbers of ions and water molecules in the bulk solution. The simulations show that, except for positively charged residues, the preferential interaction coefficients are lower for glutamate than chloride (Fig. 6b). This implies that glutamate is preferentially excluded from peptide sites. Preferential exclusion of glutamate will contribute to the increased association of macromolecules, and this explains the increased clustering in glutamate when compared to chloride ^20, 21, 22^. For Arg and Lys, the anions localize around the amino acids to neutralize the charge. As a result, we do not observe substantial differences in the preferential interaction coefficients of glutamate versus chloride around Arg and Lys, respectively.

**Figure 6:**
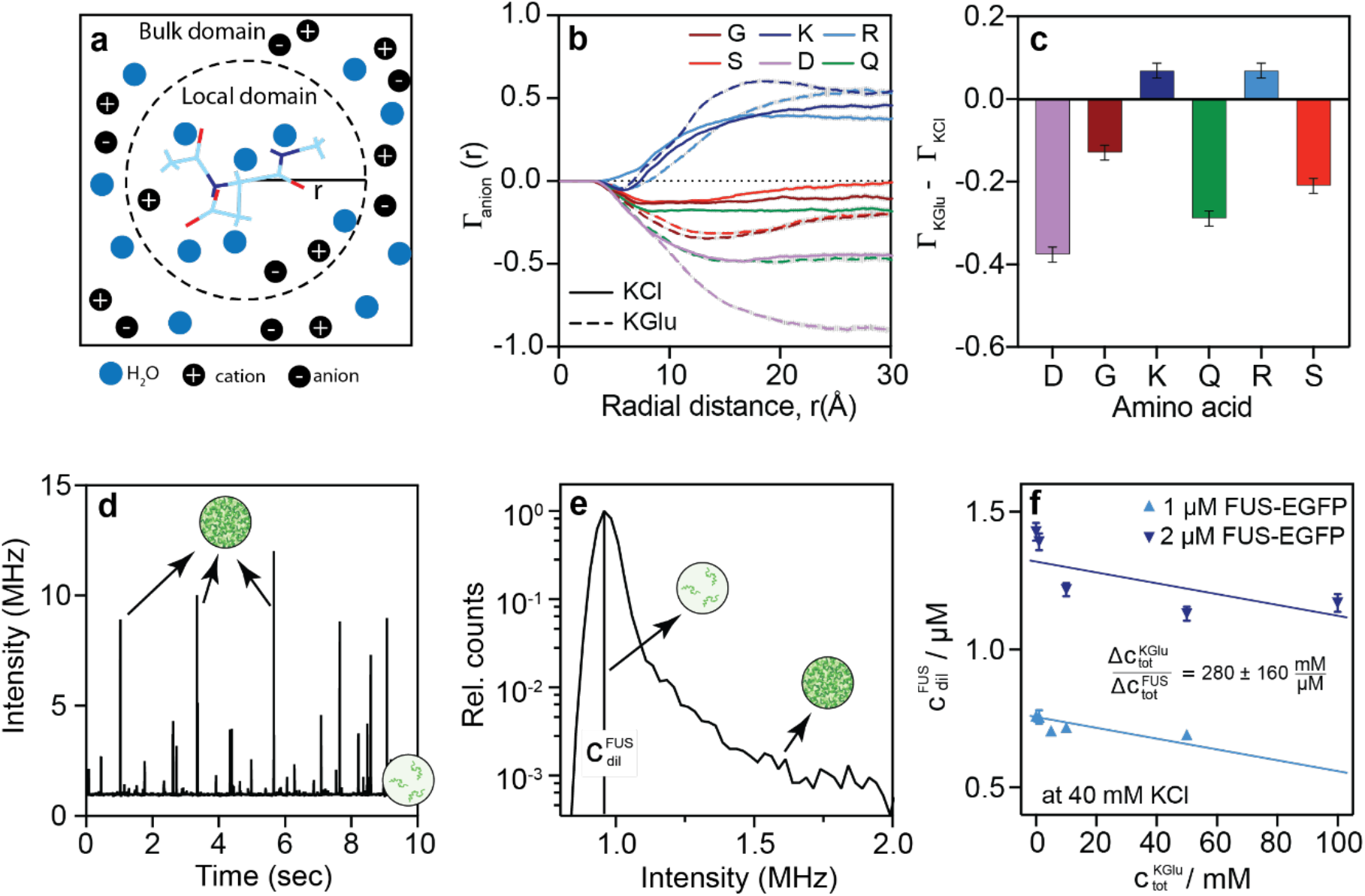
Preferential exclusion of glutamate from peptide amide sites drives protein associations and increased partitioning into dense phases. (a) Schematic for the calculation of preferential interaction coefficients as a function of distance *r* from the center-of-mass of the peptide of interest. (b) Preferential interaction coefficients for different amino acids as a function radial distance *r.* The computations are of the anion-specific preferential interaction coefficients in 500 mM KCl and 500 mM KGlu, respectively. Error bars denote the standard error of the mean. (c) The difference in the preferential interaction coefficient for KGlu and KCl (Γ_KGlu_ − Γ_KCl_) at *r* ≈ 30 Å for different amino acids. (d) Representative intensity time trace of FUS-EGFP samples post induction of phase separation using a single photon counting detection unit. (e) The intensity histogram of the time trace is shown in (d). (f) The *c*sat of FUS-EGFP decreases when KGlu concentrations are increased at a constant KCl concentration of 40 mM.

Next, we computed the preferential interaction coefficients for the whole salts using Γ_salt_ = 0.5(Γ_01231_ + Γ_cation_ − |Z|), where Γ_anion_ and Γ_cation_ are the preferential interaction coefficients for the anion and cation respectively, and Z is the net charge of the peptide (Fig. 6c). As with the values of Γ_anion_, we find that the preferential interaction coefficients are smaller for KGlu when compared to KCl. This is true for all non-cationic amino acids. Radial distribution functions between the solution anions and different moieties on different peptides (**Extended Data** Figs. 5-8) show that the differences between glutamate and chloride originate from the differences in accumulation versus exclusion of the anions around atoms of the backbone and sidechain groups.

### Preferential exclusion of glutamate enhances partitioning of FUS into dense phases

We quantified the *c*_sat_ of FUS-EGFP using a single photon counting confocal detection unit connected to a microfluidic flow cell. This allows for the recording of intensity time traces of clusters characterized by short bursts in intensity stemming from condensates passing through the confocal volume as well as the formation of a stable baseline intensity corresponding to the FUS-EGFP dilute phase concentration (Fig. 6d.). This can then be extracted from the maximum intensity histogram as the dilute phase volume fraction far exceeds that of the dense phase (Fig. 6e). These measurements were repeated for varying KGlu concentrations at two different concentrations of FUS-EGFP to record how the *c*_sat_ changes with glutamate concentrations (Fig. 6f). Here, the background KCl concentration was kept constant at 40 mM. The *c*_sat_ of FUS-EGFP decreases upon increasing KGlu. Taken together with the analysis of preferential interaction coefficients, we interpret this result as a lowering of the dilute phase concentration of FUS-EGFP due to the preferential exclusion of glutamate from protein sites. As a result of linkage thermodynamics ^36^, this enhances the free energy of transfer of proteins, including dilute-phase clusters, into the dense phase, resulting in increased partitioning of FUS-EGFP into dense phases as glutamate concentrations increase.

## Discussion

In this work, we investigated the effects of glutamate versus chloride on both macrophase separation and clustering in sub-saturated solutions of FUS and specific FET family proteins. On the molecular scale, we find that glutamate is preferentially excluded from amides and non-cationic groups when compared to chloride ^18, 19, 20, 21, 22, 35, 37, 38^. Preferential exclusion enhances protein-protein interactions such that for a given protein concentration, clustering in sub-saturated solutions is enhanced in glutamate when compared to chloride. Unlike the effects on clustering in sub-saturated solutions, the driving forces for macrophase separation, as quantified by *c*_sat_, are minimally affected by 100 mM KGlu versus 100 mM KCl.

The totality of results suggests the following: The molecular driving forces for clustering in sub-saturated solutions and phase separation may appear to be similar, but they are in fact distinct and separable. This was realized through comparative studies of the effects of glutamate versus chloride. Clustering is significantly enhanced in the presence of glutamate when compared to chloride. Preferential exclusion of glutamate enhances molecular associations. The differences in interactions between glutamate and chloride prevail on length scales of ∼1 nm. The multiplicative effect of the differences in glutamate versus chloride are manifest at ultra-low concentrations, and they are amplified as concentrations increase. However, on length scales where mesoscale clusters form and coalesce to form condensates, the differences in interactions in glutamate versus chloride become negligible. This is manifest in the minimal changes to *c*_sat_, which is the concentration at which the chemical potentials of macromolecules and solvent components are equalized across the dilute and dense phases.

The existence of a saturation concentration comes from an instability in the free energy of mixing. The implication of similar *c*_sat_ values is that the instability in the free energy of mixing, which determines the effective solubility is essentially the same in glutamate versus chloride. This further implies that the renormalized Flory chi parameter ^39^ is essentially the same in chloride versus glutamate. In the mean-field formalism of Flory ^40^ and Huggins ^41^, χ is proportional to the algebraic difference between the effective protein-solvent interactions and the sum of the protein-protein and protein-solvent interactions ^3^. Preferential exclusion of the solvent enhances protein-protein interactions on the molecular scale. Clustering does not lead to an instability in the free energy of mixing ^25^. Based on minimal changes in *c*_sat_ we infer minimal changes in the renormalized χ that determines the onset of instability in the free energy of mixing ^3, 39^. This suggests that preferential exclusion, which enhances protein-protein interactions on the molecular scale, must be counterbalanced by weaker solvent-solvent interactions through. As a result, cluster formation in sub-saturated solutions is enhanced, but *c*_sat_ changes only minimally.

*In vivo*, a clear illustration of the separation of scales and the differences between clusters in sub-saturated solutions versus condensates in supersaturated solutions comes from the recent work of Lan et al., on the nuclear protein negative elongation factor (NELF) ^8^. These proteins form condensates in response to stress, thereby arresting transcription ^42^. Using highly inclined and laminated optical sheet microscopy, Lan et al., discovered that NELF proteins in unstressed cells form heterogeneous distributions of clusters in the nucleus ^8^. The size distribution of clusters is heavy-tailed. Stress, in the form of heat-shock, drives clusters to coalesce and form condensates that sequester essential transcription factors. The separation of energy scales was discerned by quantifying responses to the MAP kinase P38. Inhibition of this kinase also inhibits condensate formation, although the clusters still persist. This suggests that multisite phosphorylation of the NELFs provide the additional energy scale that enables coalescence of clusters that drives condensate formation of NELFs. Given that these clusters form in cellular milieus, we presume that they are a reflection of contributions of anions such as glutamate.

Glutamate is an abundant metabolite in bacterial and eukaryotic intracellular milieus ^13, 14, 18, 19, 20, 21, 22, 24, 43, 44, 45, 46, 47^. The separation of energy scales and the enhanced clustering in sub-saturated solutions has likely implications for the extent of clustering at low protein concentrations in cells. The upshot is that even if the endogeneous levels of FUS and FET family proteins are below *c*_sat_, we should expect that the unbound proteins form heterogeneous distributions of clusters in cellular milieus. The abundance of clusters and the size distributions of clusters will be governed by the expression levels. These clusters, which are different from condensates, may be functionally relevant, especially in the context of transcriptional regulation ^48, 49, 50, 51^.

In neurons, cytoplasmic glutamate is concentrated in synaptic vesicles by vesicular glutamate transporters, and proton / glutamate antiport proteins ^52^. Abnormal glutamate metabolism has been reported in the context of amyotrophic lateral sclerosis (ALS) ^44, 47^. Specifically, glutamate levels have been found to increase significantly in neurons of early-stage ALS patients ^44^. However, how these elevations of glutamate cause pathological inclusions in ALS/ FTD remains unknown. The present study suggests a plausible connection between glutamate and FET proteins, which are also associated with ALS ^53^. We propose that the higher concentrations of glutamate in motor neurons of patients with ALS might result in aberrant clustering within sub-saturated solutions. We have shown that fixed concentrations of 100 mM KGlu enhances cluster formation. However, in neurons, the concentrations of glutamate vary across the cell body (∼5 mM), axon (∼10-30 mM), and synaptic vesicles (∼100 mM) ^14^. In physiological conditions, cluster formation and their abundance are likely to be influenced by concentration differences of glutamate across the neurons.

Overall, our results show that the condensate formation in supersaturated solutions and clustering in sub-saturated are governed by separable energy scales. The separation of scales is masked in chloride and unmasked in the presence of glutamate. Our work highlights how components of the cellular milieu contribute to amplifying the distinct contributions from associative versus segregative phase transitions namely, pre-percolation / percolation and phase separation, respectively ^3^. Our findings emphasize the need to account for the totality of processes that are involved in driving the formation of different types of assemblies of multivalent proteins.

## Methods

### Protein purification

All proteins were expressed in 1 L SF9 (1 million/mL) insect cells with 5 mL P2 virus and harvested 72 hours post-infection. Cells were collected by centrifugation for 5 min at 2,000 rpm. The pellets were resuspended in a lysis buffer. Protease inhibitors were added to the lysis buffer solution. The cells were lysed by sonication. The crude lysate was clarified by centrifugation for 20 min at 17,000 rpm. After centrifugation, the supernatant was passed through with a Ni-NTA agarose column using Ismatec peristaltic pump. The protein-bound beads inside the column were further washed with 10 column volumes of lysis buffer and the mixture of 10 mM of Imidazole with lysis buffer, respectively, to remove non-specific bound proteins from the Ni-NTA column. The proteins were eluted with the NTA elution buffer.

The eluted protein solution was passed through MBP resins using a gravimetric column. The protein-bound beads were washed with 10 column volume of lysis buffer. The proteins were eluted with the MBP elution buffer. The entire process was monitored using a Bradford assay, and each fraction was investigated with gel electrophoreses.

To cleave the His-MBP tag, 3C precision protease was added to the eluted protein at a 1:100 ratio. The mixture was incubated at room temperature for 4 hours and was purified using gel filtration chromatography (ÄKTA with Superdex-200 increase column; GE Healthcare) equilibrated with storage buffer. Peak fractions were pooled and used immediately for the DLS or NTA experiments. To obtain untagged protein, TEV protease was added at a 1:50 ratio and incubated at room temperature for six hours. The untagged protein was purified using gel filtration chromatography (ÄKTA with Superdex-200 increase 10/300 column; GE Healthcare) and equilibrated with storage buffer. Peak fractions were pooled and concentrated using Amicon 15 30,000 MWCO at 4000 rpm at room temperature. Protein concentrations were determined by measuring absorbance at 280 nm using a NanoDrop ND-1000 spectrophotometer (Thermo Scientific). The 260/280 ratio of all purified proteins was measured between 0.52 to 0.56. Proteins were stored with tags at −80°C, and prior to experiments, the proteins were thawed, the tag was cleaved, gel filtration chromatography was run, and peak fractions were pooled, concentrated, and used for the experiments. This protocol allows for data reproducibility. Details of all the constructs used in the experiments are described in the *Supplementary Information*.

### Determination of saturation concentrations

We prepared the samples at the indicated protein concentration in 100 mM KCl and KGlu buffers, and 30 minutes after sample preparation, the samples were spun down at 20,000 RCF on a benchtop centrifuge unit and measured the concentration in the clarified supernatant by using a Bradford assay ^54^.

### Microscopy measurements of phase separation

For droplet formation assays, proteins were diluted into various concentrations in the corresponding 100 mM KCl buffer, 100 mM KGlu buffer, and KCl-control buffers in a total solution volume of 30 μL. The samples were added into the 384 well non-binding microplates (Greiner bio-one). The images were taken after various time points, starting from 1 hour to 4 hours. Images were taken using an IX71/IX81 inverted Spinning Disc Microscopes with an Andor Neo sCMOS/Andor Clara CCD camera and an UPlanSApo 60x oil-immersion objective (Olympus).

### DLS Measurements

DLS measurements were performed using the Zetasizer Nano ZSP Malvern instrument (measurement range of 0.4 nm to 10 µm). The Nano ZSP instrument incorporates noninvasive backscattering technology. This enables the measurement of time-dependent fluctuations of the intensity of scattered light as scatterers undergo Brownian motion. The analysis of these intensity fluctuations enables the determination of the diffusion coefficients of particles, which are converted into a size distribution using the Stokes-Einstein equation ^55^. A 632.8 nm laser illuminated the sample solutions, and the intensity of light scattered at an angle of 173° was measured using a photodiode.

In DLS, the autocorrelation function of the scattered light is used to extract the size distribution of the dissolved particles. The first order electric field correlation function of laser light scattered by a monomodal or monodisperse population of macromolecules can be written as a single exponential of the form:

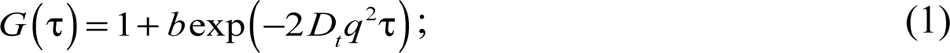

Here, *b* is a constant that is determined by the optics and geometry of the instrument, *D_t_* is the translational diffusion coefficient of the particles, and τ is the characteristic decay time. The scattering vector *q* is given by:

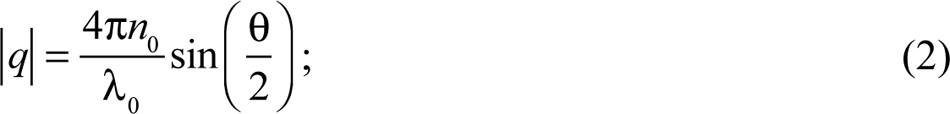

Here, *n*_0_ is the refractive index of the solvent, λ_0_ is the wavelength, and θ is the scattering angle. For populations composed of a single type of scatterer, the distribution function of decay rates can be derived from a simple fit of the experimental estimates of the logarithm of the correlation function in Eq. 1 to a polynomial. These methods, which apply to monomodal distributions of sizes of scatterers, can be used to extract the translational diffusion coefficient, from which one can estimate the hydrodynamic radius *R_h_* of the scatterers. For this, one uses the Stokes-Einstein relation:

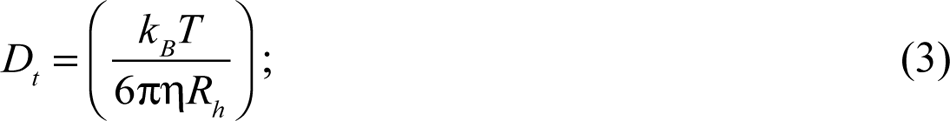

Here, *k_B_* is the Boltzmann constant (1.381 × 10^-23^J/K) and η is the absolute (or dynamic) viscosity of the solvent. In this work, we used the hydrodynamic diameter *d_h_* (i.e., *d_h_*= 2*R_h_*) as the preferred way to quantify particle sizes.

For the measurements, all solutions were filtered using 0.2 μm membranes (Millex®-GS units) purchased from Millipore™. All experiments were conducted with the following settings on the Malvern instrument: Material – protein; Dispersant – buffers; Mark-Houwnik parameters; Temperature: 25 °C with equilibration time – 120 seconds, Measurement angle: 173°. Each spectrum represents the average of 12 scans every 10 seconds in duration. All proteins were freshly purified and used after chromatography purification with a standard stock solution buffer. The samples were prepared by adding freshly prepared stock proteins followed by dilution buffer and mixed thoroughly by pipetting 4 to 6 times. The samples were equilibrated for 2 mins at 25 °C, and the data were recorded in 2-minute intervals.

### Nanoparticle Tracking Analysis (NTA)

Nanoparticle tracking analysis was performed using NS300 from Malvern instruments (measurement range of 20 nm to 1µm). The system was accompanied by a NanoSight syringe pump to inject the samples for the experiments. NTA measurements utilize the properties of light scattering and Brownian motion to obtain the size distributions and concentrations of particles in liquid suspension. A laser beam (488 nm) was passed through the sample chamber, and the particles in suspension were visualized using a 20x magnification microscope. The video file of particles moving under Brownian motion was captured using a camera mounted on the microscope that operates at 30 frames per second. The software tracks particles individually and uses the Stokes-Einstein equation to resolve particles based on their hydrodynamic diameters.

All proteins were freshly purified and used after chromatographic purification with a standard stock solution buffer. All buffers were filtered through a 0.22 μm polyvinylidene fluoride membrane filter (Merck, Germany). All protein stock solutions were centrifuged at 20000 RCF for 5 mins at room temperature before measurements. The samples were prepared by adding freshly prepared, centrifuged stock proteins followed by dilution buffer and mixed thoroughly by pipetting 4 to 6 times. The samples were equilibrated for 2 minutes at 25°C, and the data were recorded 6 minutes after sample preparation and equilibration.

### Multiparameter fluorescence detection (MFD) measurements and analysis

#### Confocal point measurements

All confocal point measurements were conducted on a confocal fluorescence microscope (Olympus IX71, Hamburg, Germany) using a supercontinuum laser (SuperK Extreme with SuperK VARIA tunable filter, NKT Photonics, Birkerød, Denmark) at 514 nm ± 1.5 nm and 39 MHz. Laser light was directed into a 60x water immersion objective (NA = 1.2) by a dichroic beam splitter and focused into the sample close to the diffraction limit volume. The emitted light was collected by the same objective and separated into two polarizations (parallel and perpendicular) relative to the excitation beam. The fluorescence signal was further divided into two spectral ranges by beam splitters (BS 560, AHF, Tübingen, Germany). Additionally, Bandpass filters for Nile Red fluorescence (HC 607/70) were placed in front of the detectors. The signals from single photon counting detectors (SPCM-AQRH-14 TR Excelitas, Wiesbaden, Germany) were recorded photon-by-photon with picosecond accuracy (HydraHarp400, PicoQuant, Berlin, Germany) and analyzed using custom software (LabVIEW based, see https://www.mpc.hhu.de/software/mfd-fcs-and-mfis). The temperature in the laboratory during all titration steps was 20 ± 1 °C.

FUS-SNAP or untagged FUS was titrated into 15 nM Nile Red solution at standard buffer conditions (20 mM TRIS, pH 7.6) with either 100 mM KCl or 100 mM KGlu. The KGlu buffer contains 10 mM residual KCl from the FUS stock solution. Both dye and salt concentrations are kept constant during the titration and back-dilution using the respective Nile Red-containing buffer solutions.

#### Burst analysis

Using in-house developed software (LabVIEW based) bursts were selected via an intensity threshold using a 2σ criteria out of the mean background with a photon minimum number of 10 photons per burst. The decays for each burst were then processed and fitted using a mono-exponential model yielding fluorescence-weighted average lifetime considering correction factors for background signal, polarization (g-factor) and scattering effects. Especially for FUS titrations in the nM to µM regime, an additional photon maximum of 3000 was applied due to computational limitations.

#### Fluorescence intensity distribution analysis (FIDA)

The FIDA analysis for all experiments was executed according to the established protocols ^56^.

##### Single-molecule FRET (smFRET) measurements

smFRET experiments were conducted using a confocal epi-illuminated setup based on an Olympus IX71 microscope deploying pulsed interleaved excitation (PIE) where the donor and acceptor fluorophore are sequentially excited by fast alternating laser pulses thus allowing the computation of the stoichiometry S (donor-acceptor-ratio). Excitation is attained using 485 nm (50 μW) and 640 nm (10 µW) pulsed diode lasers (LDH-D-C 485 and LDH-P-C-635B, PicoQuant Berlin, Germany) operated at 32 MHz and focused by a 60×/1.2 NA water immersion objective (UPLAPO 60x, Olympus, Hamburg, Germany) into the sample. We used the excitation beam splitter FF500/646 (Semrock, USA), a polarizing beam splitter cube (VISHT11, Gsänger) and dichroic detection beam splitters (595 LPXR, AHF, Tübingen, Germany) to separate fluorescence from laser excitation and split it into its parallel and perpendicular spectral components. The four-detection channel, corresponding to color and polarization, were split further by 50/50 beam splitters to obtain dead time-free species cross-correlation curves, yielding a total of eight fluorescence detection channels (green channels: τ-SPAD-100, PicoQuant; red channels: SPCM-AQR-14, Excelitas, Wiesbaden, Germany). To block out Raman scattering green (HQ 530/43 nm for FUS-SNAP-Alexa488) and red (HQ 720/150 nm for FUS-SNAP-Alexa647) bandpass filters (AHF, Germany) were put in front of the corresponding detectors. The detector outputs were recorded by a TCSPC module (HydraHarp 400, PicoQuant). Measurement times for single-molecule detection (SMD) experiments were about 10 hours each ^57, 58^.

The equations for the static (S.4) and dynamic (S.5) FRET lines including the weighted lifetimes for both species, the correction parameters and all 2D-FRET efficiency plots are listed in the following.

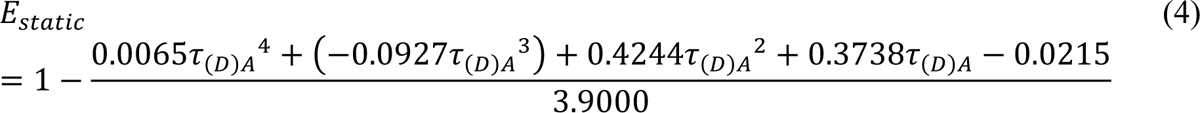

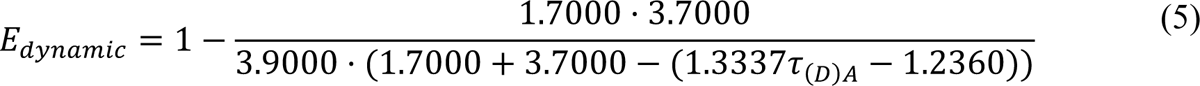

### Fluorescence correlation spectroscopy (FCS) curve fitting

Donor autocorrelations of smFRET measurements were fitted for correlation times *t_c_* from 10^-4^ to 10^2^ ms using a 3D-Gaussian model with two diffusion terms, corresponding to a dye/monomer component *t_d1,global_* and a multimer component *t_d2_*. The parameters in Equation (S.6) are tabulated in the *Supplementary Information*.

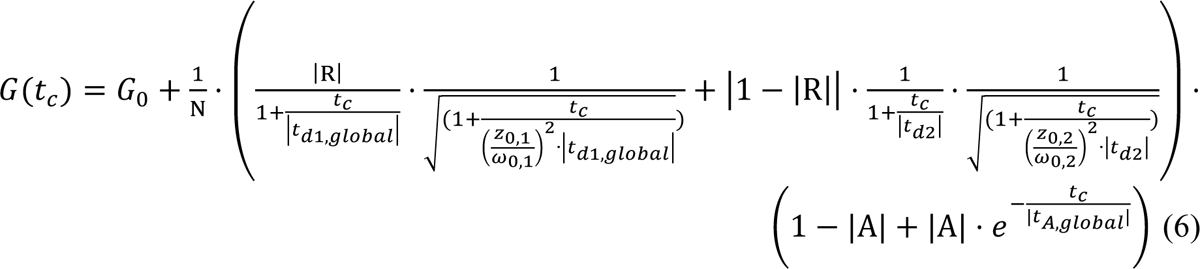

### Microfluidic Confocal Spectroscopy (MCS)

Microfluidic devices ^27^, were first fabricated as SU-8 molds (MicroChem) through standard photolithographic processes and then produced as polydimethylsiloxane (PDMS) slabs, which were bonded onto thin glass coverslips ^59^. The devices were operated by placing gel-loading tips filled with buffer and protein sample in their corresponding inlet ports and pulling solution through the devices in withdraw mode at a flow rate of 150 µL/h using automated syringe pumps (neMESYS, Cetoni).

The FUS-EGFP was stored in 500 mM KCl, 20 mM TRIS.HCl pH 7.4, was diluted with buffers of 20 mM TRIS.Glu to the indicated protein and KGlu concentrations as stated. During the experiment, the sample was placed into the sample inlet of the device, and the corresponding buffer contained the same concentration of KGlu in the buffer inlet. The co-flowing buffer was supplemented with 0.05% Tween-20 to prevent surface sticking of the protein to PDMS and glass surfaces.

Experiments were conducted by scanning the confocal spot of a custom-built confocal microscope through the central four channels of the microfluidic device ^7^. Briefly, the setup is equipped with a 488-nm laser line (Cobolt 06-MLD) for excitation of EGFP fluorophores and a single-photon counting avalanche photo diode (SPCM-14, PerkinElmer) for subsequent detection of emitted fluorescence photons. Further details of the optical unit have been described previously. During the scanning of the device, 200 evenly spaced locations within the central four channels of the device were surveyed and detected for 4 seconds. Clusters were classified as peaks that exceeded 5 standard deviations above the mean fluorescence intensity of each trace. These peaks were quantified according to location against the mean signal of each trace. The average number of clusters *n*_clusters_ was then quantified by averaging each of the four groups of peaks corresponding to the four central channels. This was used in the calculation of cluster concentration according to the following equation:

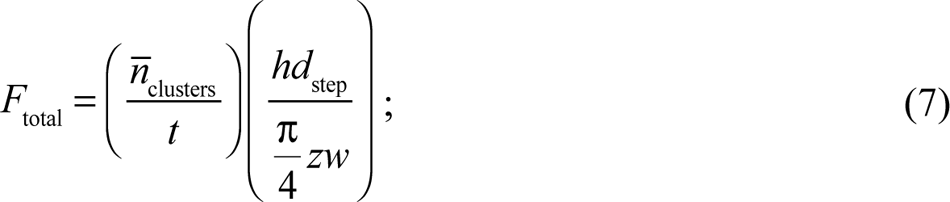

Here, *t* is the time each trace was collected for (4 seconds), *h* is the height of the microfluidic channel (28 μm), *d*_step_ is the width of each step (5.64 μm), and *z* and *w* were the height and width of the confocal spot (3 μm and 0.4 μm, respectively). From Equation (S.7), which yields the flux of clusters *F*_total_, the concentration of clusters could be determined according to Equation (S.8), with *Q*_sample_ being the flow rate of the sample (15 μL/h) and *N*_A_ being the Avogadro constant ^29^:

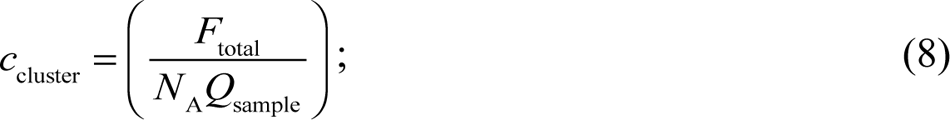

### Bis-ANS experiment

Different concentrations of FUS-SNAP protein were mixed with 2 μM bis-ANS solution in 100 mM KCl and KGlu buffers. Then the solutions were mixed and loaded 100 µL in a 96 well plate (microplate, PS, half area, µClear, Med. binding, Black, Greiner Bio-one). The spectra were recorded from 425 nm to 650nm (10 nm bandwidth) with the TECAN plate reader using an excitation wavelength of 355±5 nm. For control, we used only 2 μM bis-ANS with the same buffer conditions, ethanol, and methanol.

### Molecular Dynamics (MD) simulations

We used Charmm36 ^60^ forcefield to perform MD simulations using Gromacs 2021 package ^61, 62^. Simulations were performed with explicit representations of solvent molecules using the TIP3P^63^ water model. The simulation set up was as follows: we used capped amino acids of the form Ace-Xaa-Nme, where Ace refers to N-acetyl, Nme refers to N′-methylamide, and Xaa is the residue of interest, which is one of Gly, Asp, Glu, Arg, Lys, or Gln. Then, we solvated the peptide by using the default “scale” factor of 0.57 in a cubic simulation box with box size 7 × 7 × 7 *nm*^3^. Ions were added to the simulation box, by replacing water molecules to neutralize the charge on amino acid (if any) and to obtain salt concentration of 500 mg / mL. On average, we obtained 11,007 water molecules for the simulations where KCl is used and 10,333 water molecules for simulations with KGlu.

We then performed energy minimization using the steepest descent method followed by 100 ns equilibration at 298K and 1 bar. We then performed additional simulations, each 400 ns long to obtain production runs which were later used to obtain radial distribution functions and preferential interaction coefficients. Periodic boundary conditions were employed in all three directions. The V-rescale thermostat^64^ was used to maintain the temperature at 298 K with a coupling time constant of 0.1 ps. The pressure was maintained by using the Parinello-Rahman^65^ method with a time constant of 2.0 ps. Long-range electrostatic interactions were handled using the smooth Particle-mesh Ewald (SPME) algorithm. We used the cutoff for short-ranged Lennard-Jones potential to be 1.1 nm while the cutoff for the short-ranged electrostatic potential was 1.2 nm. The LINCS algorithm^66^ was used to constrain covalent bonds involving hydrogen atoms.

#### Calculation of preferential interaction coefficients

Preferential interaction coefficient is a measure of the amount of co-solute in the local domain of the peptide compared to the solvent. We determined the number of ions (*N*_8MC5(N)_) and the number of water molecules (*N*_H2O_(γ)) as a function of distance γ from the center of mass of the peptide, by using distance bins of size 0.2 Å. We then found the ratio of bulk density of ions to water molecules by calculating the ratio 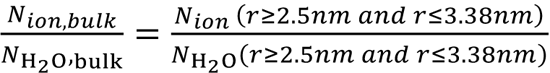 We used the 400 ns of the production run across 4 replicates to obtain preferential interaction coefficients as a function of distance γ from the center of mass of the peptide.

## Supplementary Information

Additional details of methods and the analysis of results can be found in the *Supplementary Information*.

## Acknowledgments

We thank Furqan Dar, Tyler Harmon, Ralf Kühnemuth, Min Kyung Shinn, and Jie Wang for helpful discussions and assistance. This work was funded by a direct grant from the Max Planck Society (to AAH), a grant from the NOMIS foundation (to AAH), the Wellcome trust (209194/Z/17/Z to AAH), the European Research Council through the ERC grant PhysProt (to TPJK, agreement no. 337969), the Wellcome Trust and the Frances and Augustus Newman foundation (to TPJK), SPP2191 from the Deutsche Forschungsgemeinschaft (to CAMS and RVP), the US National Institutes of Health (R01NS121114 to RVP), and the St. Jude Children’s Research Hospital collaborative research consortium on membraneless organelles (to RVP). We are grateful for the technical support provided by Régis Lemaitre and Barbara Borgonovo of the protein expression, purification, and characterization facility of the MPI-CBG. We thank Andrei Pozniakovsky for the DNA constructs of all proteins.

## Author Contributions

MK, AAH, CAMS, and RVP designed the project. MK discovered the effects of glutamate on the clustering of FET family proteins in sub-saturated solutions. MK prepared all the samples and performed the NTA, DLS, *c*_sat_, and microscopy measurements with assistance from ARK. CAMS and LTV designed, and LTV performed the smFRET, FCS, and MFD measurements using samples provided by MK. LTV analyzed and interpreted all the single molecule and MFD measurements with guidance from and collaboration with CAMS. GC performed the MD simulations and analyzed the results with inputs from RVP. TJW, HA, and TPJK designed, performed, and analyzed the MCS measurements using samples provided by MK. The structure of the manuscripts and layout of figures were designed by MK, LTV, GC, and RVP. MK, LTV, and GC put together all the figures with inputs from CAMS and RVP. RVP wrote the manuscript with significant inputs from MK, LTV, and GC. AAH, CAMS, and RVP provided critical intellectual inputs. TPJK, AAH, CAMS, and RVP secured funding for the work. All authors read and edited the manuscript.

## EXTENDED DATA FIGURES FOR

**Extended Data Fig. 1:**
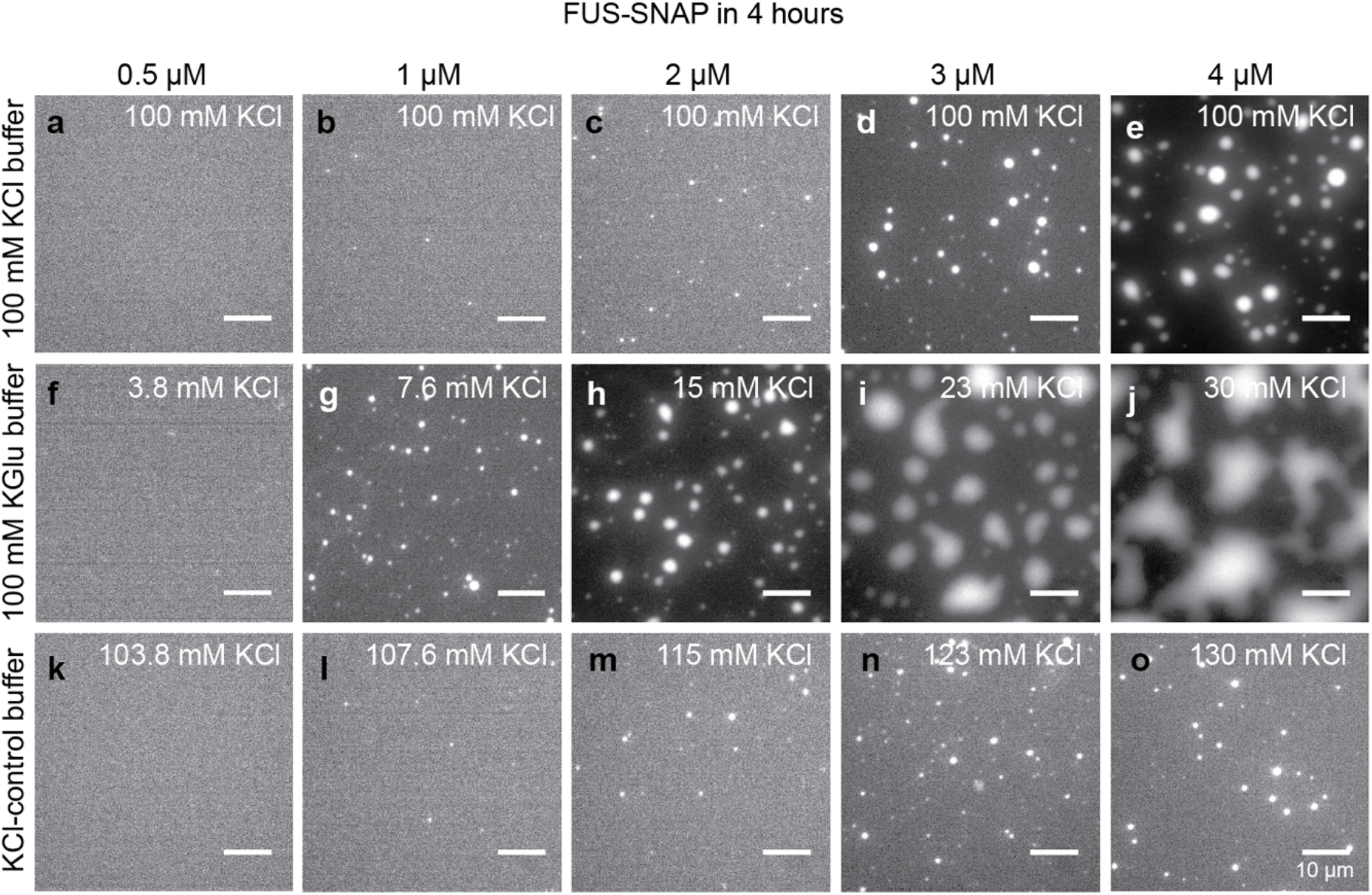
Potassium Glutamate (KGlu) buffer minimally affects the driving forces for macrophase separation of FUS-SNAP although the evolution of condensates is discernibly different. (a)-(e) shows microscopy images collected at the 4-hour time point for solutions containing different concentrations of FUS-SNAP in 20 mM Tris.HCl, pH 7.4, with a final concentration of 100 mM KCl. Panels (f)-(j) show microscopy images collected at the 4-hour time point for solutions containing different concentrations of FUS-SNAP in 20 mM TRIS.Glu, pH 7.4, with 100 mM KGlu, and panels (k)-(o) show microscopy images collected at the 4-hour time point for solutions containing different concentrations of FUS-SNAP in 20 mM Tris.HCl, pH 7.4, with 100 mM KCl. In both KGlu and KCl buffer, the residual KCl (< 30 mM) from FUS-SNAP stock was added. For imaging purposes, 5% of the total mixture in each sample is made up of Alexa-488 labelled FUS-SNAP. The total KCl concentration in the solution is marked on the panels. The scale bar in each panel corresponds to 10 µm.

**Extended Data Fig. 2:**
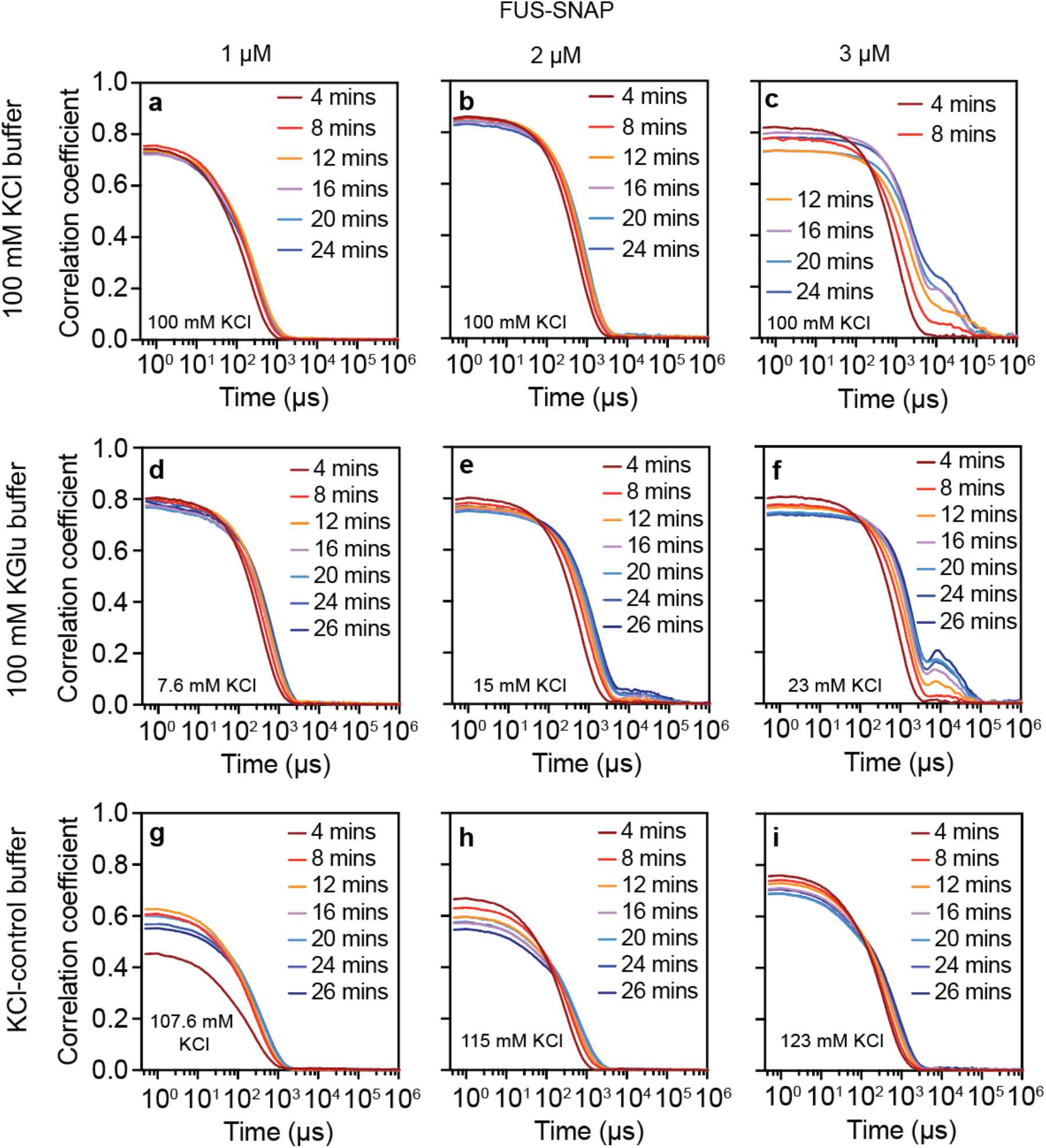
Potassium Glutamate (KGlu) buffer minimally affects the driving forces for macrophase separation of FUS-SNAP, although the evolution of condensates is discernibly different. (a)-(c) The correlation function from the dynamic light scattering of solutions containing different concentrations of FUS-SNAP, 1 μM (a), 2 μM (b), and 3 μM (c) in 20 mM Tris.HCl, pH 7.4, with a final concentration of 100 mM KCl. Panels (d), (e), and (f) show the correlation functions of solutions containing 1 μM, 2 μM, and 3 μM concentrations of FUS-SNAP, respectively, in 20 mM TRIS.Glu, pH 7.4, with 100 mM KGlu. Panels (g), (h), and (i) show the correlation functions of solutions containing 1 μM, 2 μM, and 3 μM concentrations of FUS-SNAP, respectively, in 20 mM TRIS.HCl, pH 7.4, with 100 mM KCl. The total concentration of KCl is marked on the panels. The correlation coefficient value indicates the abundance of clusters in the solutions. The time axis correlates with the size of the species, larger the size requires more time to decay and vice versa.

**Extended Data Fig. 3:**
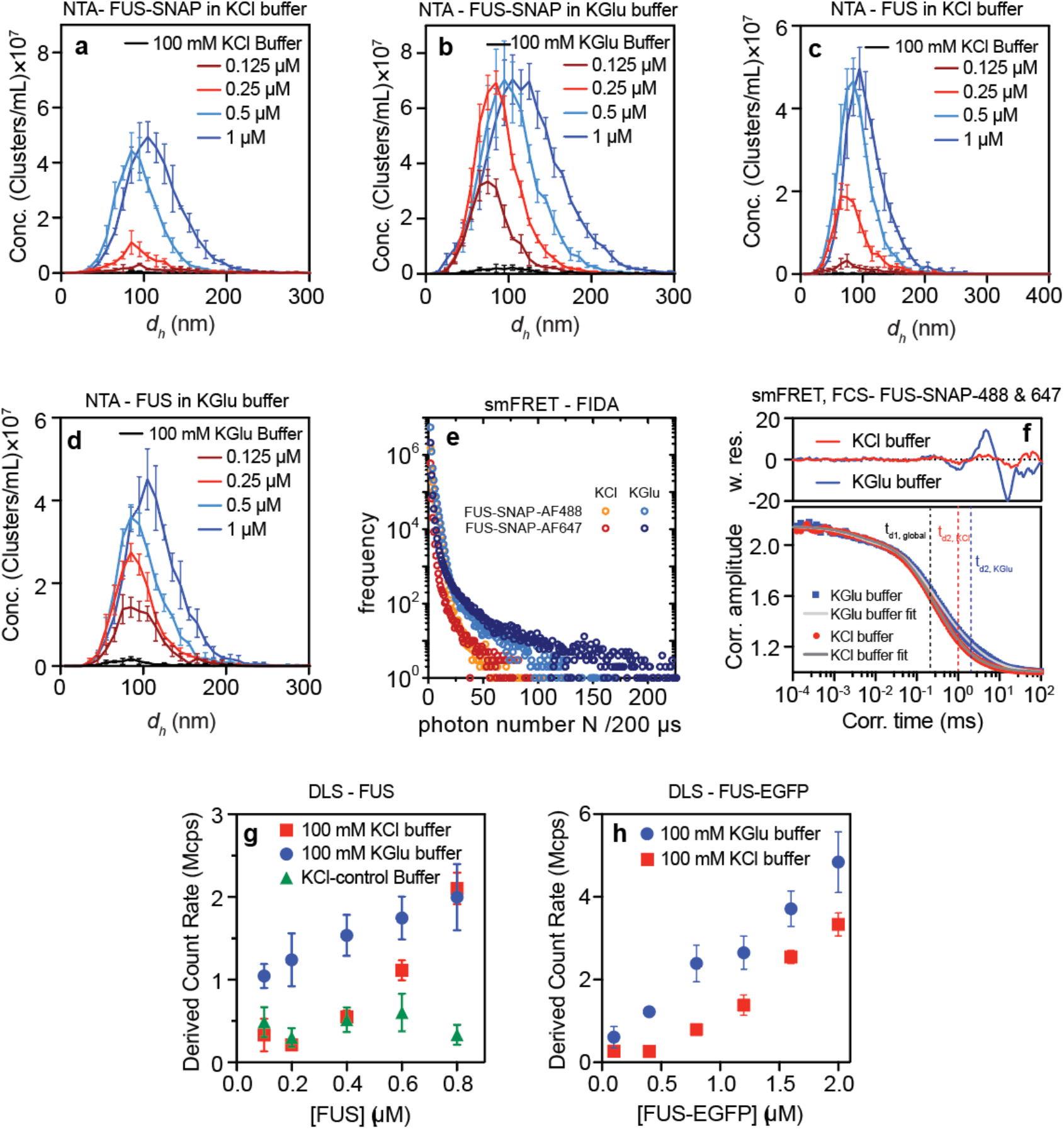
KGlu buffer enhances cluster formation of FUS-SNAP and FUS when compared to KCl-based buffer. The distribution of cluster sizes measured using NTA of (a) FUS-SNAP in 100 mM KGlu buffer, (b) FUS-SNAP in 100 mM KCl buffer, (c) FUS in 100 mM KCl buffer, and (d) FUS in 100 mM KGlu buffer (d). (e) Fluorescence intensity distribution analysis of single-molecule FRET (smFRET) measurements with 200 pM FUS-SNAP-AF488 as the donor and 200 pM FUS-SNAP-AF647 as acceptor shows more pronounced tailing towards higher photon numbers for donor/acceptor in KGlu (light blue/dark blue) than in KCl (orange/red). (f) Fluorescence correlation spectroscopy (FCS) applied to smFRET yields two translation diffusion times by fitting the autocorrelation of AF488-detection with a 3D-Gaussian diffusion model. The first diffusion time t_d1, global,_ corresponds to free dye/monomer and is, therefore, the same for both KCl and KGlu. The second diffusion time t_d2_ corresponds to FUS multimers which exhibits an almost two-fold increase for KGlu. High systematic fluctuations in the weighted residuals indicate the presence of further higher-order complexes. (g)-(h) DLS data shows the derived count rate of FUS (g) and FUS-EGFP (h) in different buffers.

**Extended Data Fig. 4:**
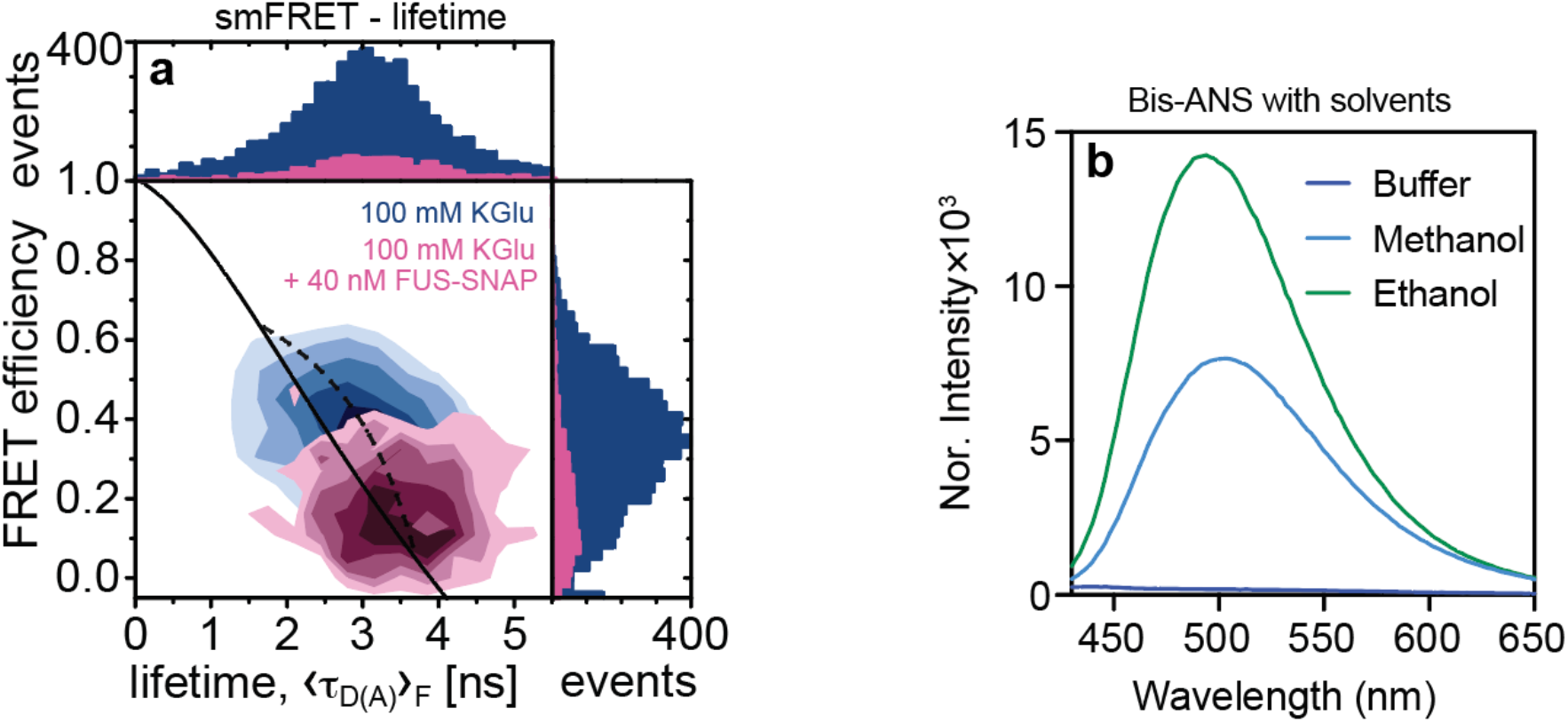
The FUS-SNAP clusters in KGlu buffer are reversible: (a) Under single-molecule conditions, adding upon the 400 pM donor (AF488) and acceptor (AF647) labelled FUS-SNAP (measurement time 10 hours) 40 nM unlabeled FUS-SNAP, we observe a significant decrease in FRET efficiency, demonstrating the reversibility of FUS association on the millisecond time scale limited by translational diffusion. (b) The control experiments were where 2 µM Bis-ANS was mixed with buffer, methanol, and Ethanol.

**Extended Data Fig. 5:**
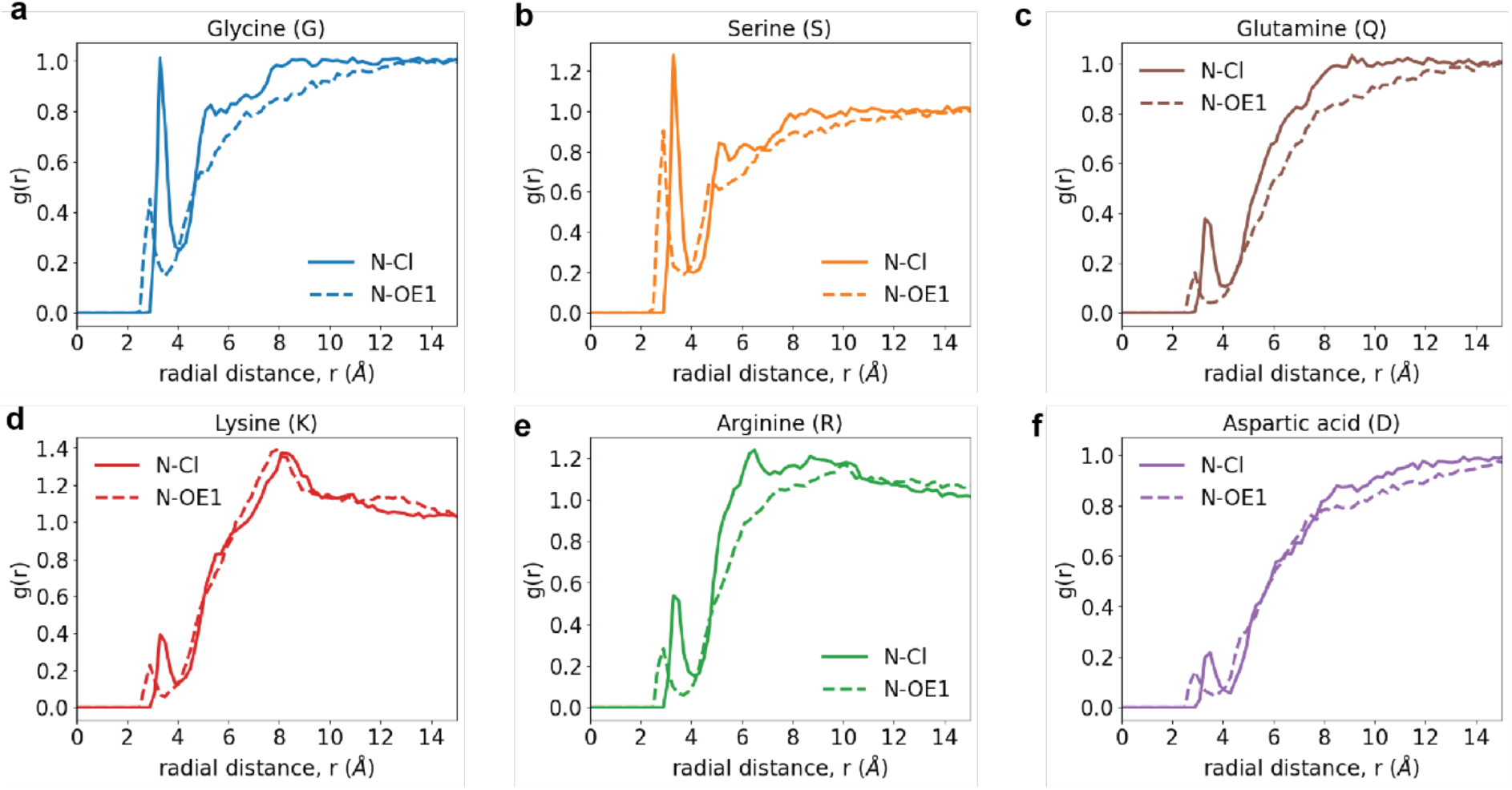
Site-site radial distribution functions, g(r) around backbone amide nitrogen atoms. These quantify the relative probability, with respect to an ideal gas prior, of finding Cl^-^ atoms (solid curve) or the *sp*^2^ oxygen OE1 of glutamate around the amide nitrogen of the backbone for different capped amino acids. Data are shown for the g(r) of Cl-(solid curve) and OE1 atom of glutamate (dashed curve) around the backbone amide nitrogen of (a) Gly, (b) Ser, (c) Gln, (d) Lys, (e) Arg, and (f) Asp. For Gly, Ser, and to a smaller extent for Gln, we observe differences in the form of lower occupancies of OE1 when compared to Cl^-^, suggesting preferential exclusion. The differences in the occupancies around the backbone nitrogen originate, in part, from the differences in sidechain length and the type of functional group that makes up the sidechain (see **Extended Data Fig. 8**). Note that the preferential exclusion extends to a spatial range of 10 – 12 Å. However, for Arg, Lys, and Asp, electrostatic considerations weaken the differences between the computed g(r) of the anions around the backbone amides. The electrostatic interactions refer to attractions of the anions to sidechains of Arg and Lys, and repulsions from the sidechain of Asp. Due to covalent connectivity, these electrostatic effects influence the occupancies around the backbone nitrogen atoms, thereby weakening the differences between the solution anions.

**Extended Data Fig. 6:**
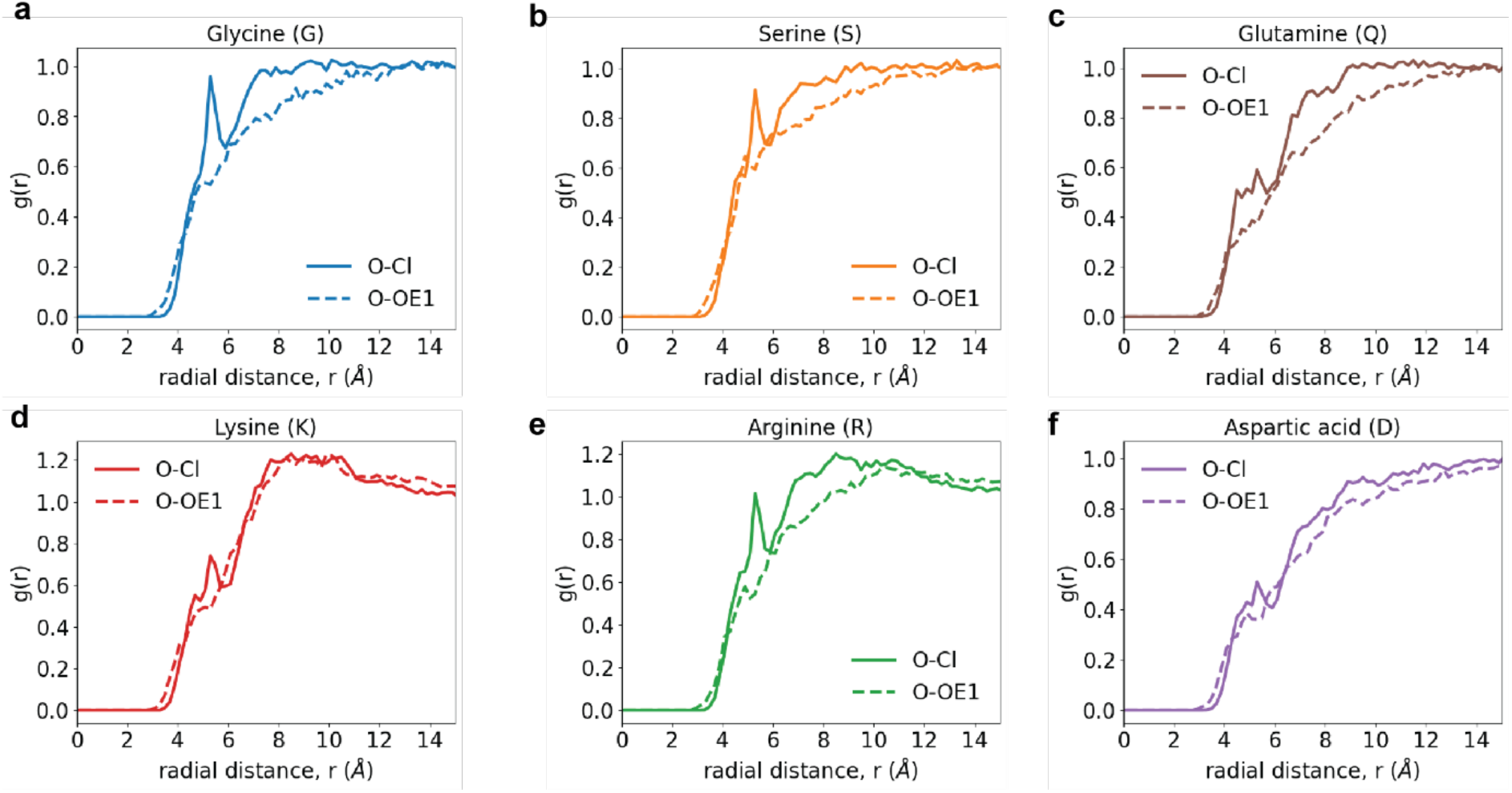
Site-site radial distribution functions, g(r) around backbone carbonyl oxygen atoms. These quantify the relative probability, with respect to an ideal gas prior, of finding Cl^-^ atoms (solid curve) or the *sp*^2^ oxygen OE1 of glutamate around the carbonyl oxygen of the backbone for different capped amino acids. Data are shown for the g(r) of Cl-(solid curve) and OE1 atom of glutamate (dashed curve) around the backbone carbonyl oxygen of (a) Gly, (b) Ser, (c) Gln, (d) Lys, (e) Arg, and (f) Asp.

**Extended Data Fig. 7:**
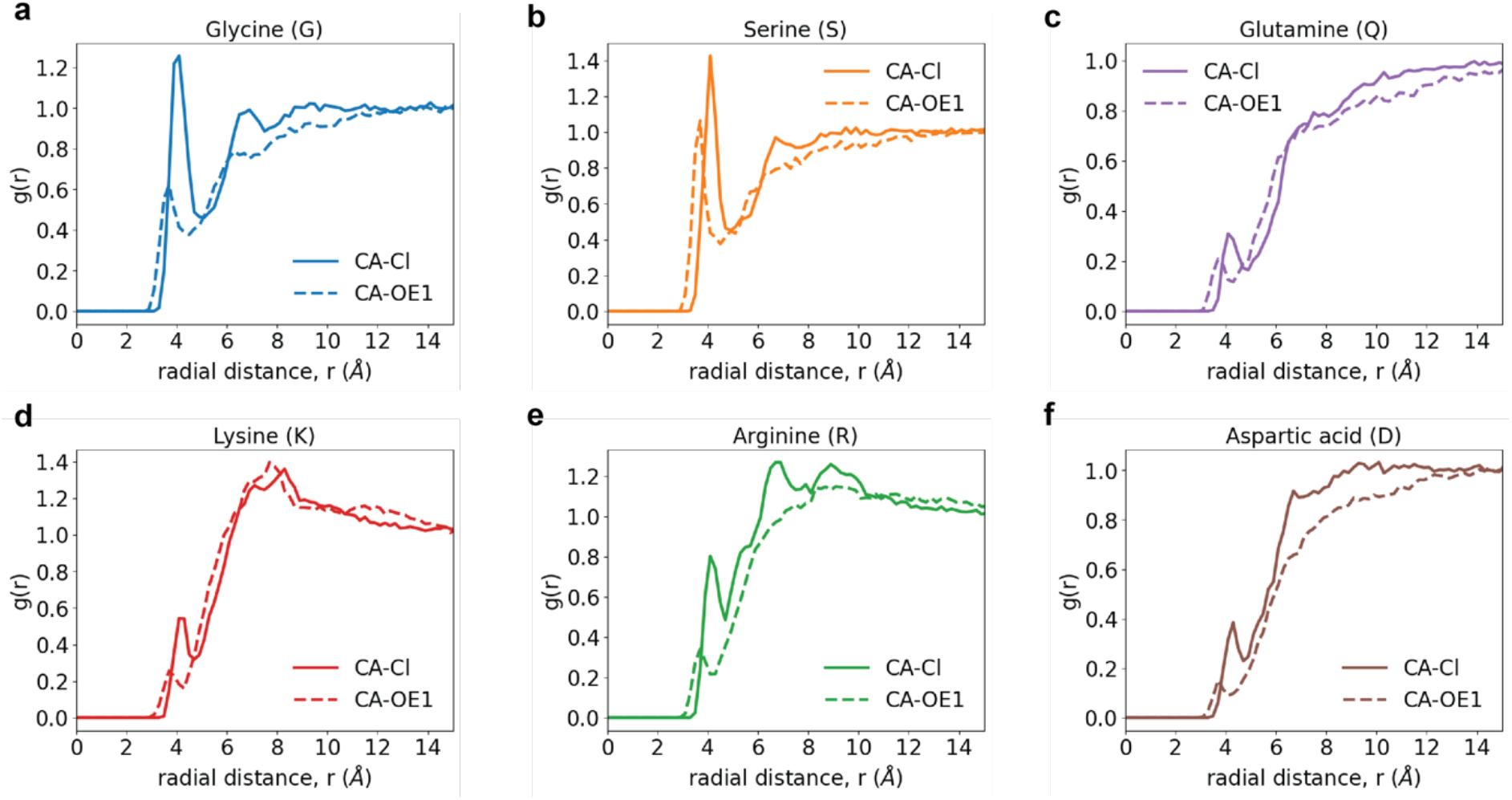
Site-site radial distribution functions, g(r) around the central (alpha) carbon atoms. These quantify the relative probability, with respect to an ideal gas prior, of finding Cl^-^ atoms (solid curve) or the *sp*^2^ oxygen OE1 of glutamate around the alpha carbon of the backbone for different capped amino acids. Data are shown for the g(r) of Cl-(solid curve) and OE1 atom of glutamate (dashed curve) around the backbone central (alpha) carbon atom of (a) Gly, (b) Ser, (c) Gln, (d) Lys, (e) Arg, and (f) Asp. While the preferential exclusion of the glutamate OE1 is pronounced around the central carbon of Gly and the alpha carbon of Ser, the differences are less pronounced around the alpha carbon of Gln. A combination of sidechain length and the site-specific strong versus weak interactions with the sidechain amides influence the occupancies around the alpha carbon atom. The influence of occupancies around the sidechain functional groups, summarized in **Extended Data Fig. 8**, influence the differences observed in the occupancies of Cl^-^ and OE1 of glutamate around the alpha carbon atoms of Lys versus Arg versus Asp.

**Extended Data Fig. 8:**
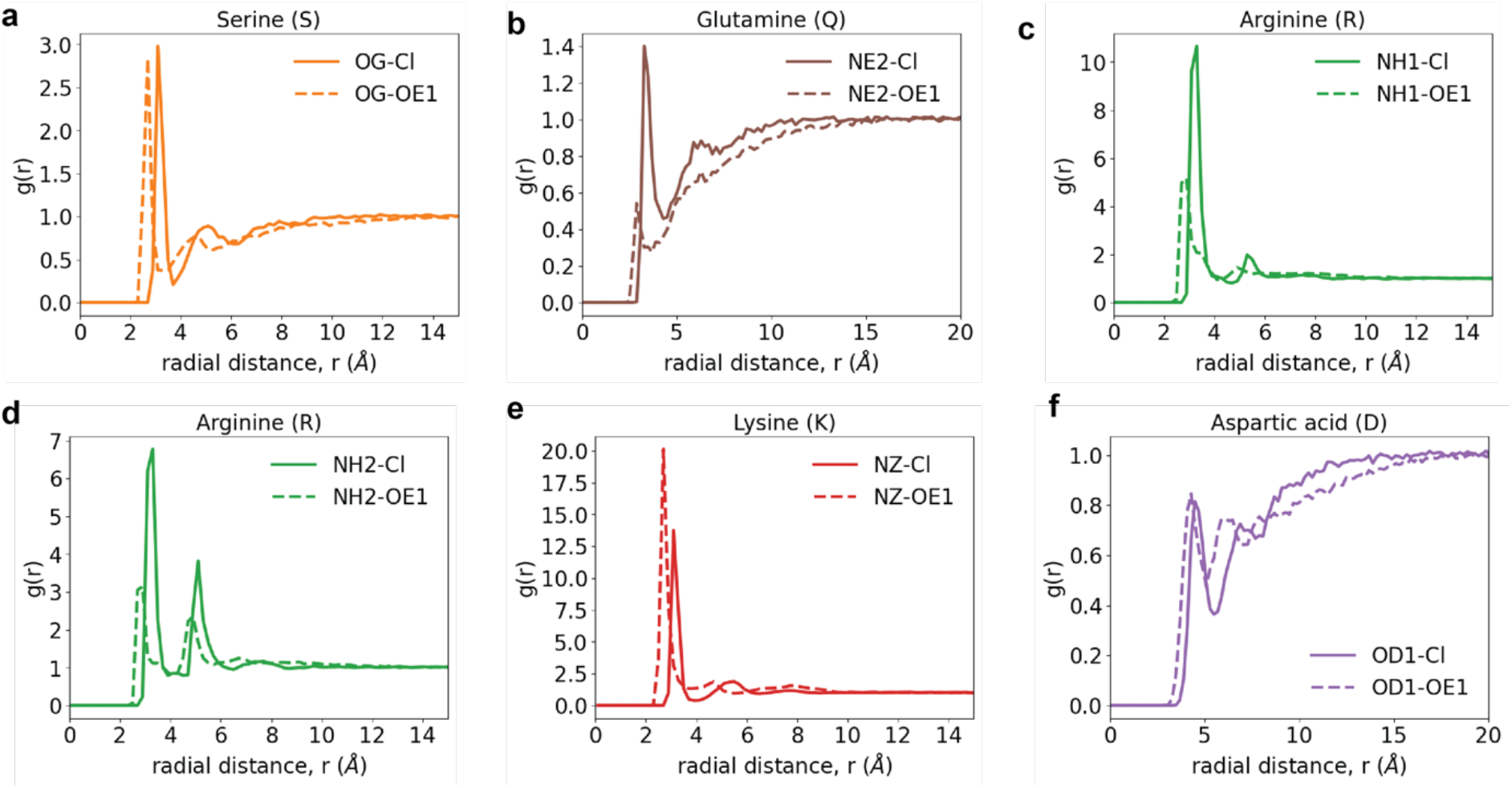
Site-site radial distribution functions, g(r) of the solution anions around specific sites of functional groups of sidechains. These quantify the relative probability, with respect to an ideal gas prior, of finding Cl^-^ atoms (solid curves) or the *sp*^2^ oxygen OE1 of glutamate (dashed curves). (a) Comparative g(r) profiles of Cl^-^ versus the epsilon 1 (OE1) oxygen of glutamate around the gamma oxygen of the hydroxyl group of Ser. The OE1 is more proximal than the Cl^-^ and this has to do with the differences in sizes of the atoms. However, the strengths of the interactions, quantified in terms of the heights of the g(r) profiles, are equivalent. Cl^-^ and the glutamate OE1 interact favorably and equivalently with the hydroxyl of Ser sidechains. (b) Comparative g(r) profiles of Cl^-^ versus the epsilon 1 (OE1) oxygen of glutamate around the epsilon 2 nitrogen, the hydrogen bond donor, of the primary amide of Gln. Again, the leftward shift of the OE1 profile has to with the smaller size vis-à-vis Cl^-^. The preferential exclusion of glutamate from amides versus the preferential interaction of Cl^-^ with these sites is evident from the comparisons of the g(r) profiles. (c) and (d) In general, the interactions of Cl^-^ are stronger with the guanido group of Arg as evidenced by the clear differences in the site-site g(r) profiles of Cl^-^ and OE1 around the eta nitrogen atoms within the guanido moieties. (e) The interactions of OE1 are modestly stronger with the amine of lysine when compared to those of Cl^-^. (f) Electrostatic repulsions contribute to roughly equivalent weakening of the interactions of both Cl^-^ and OE1 around the delta oxygen of Asp.

## SUPPLEMENTARY INFORMATION FOR

### Section B. Materials

**Table S1:**
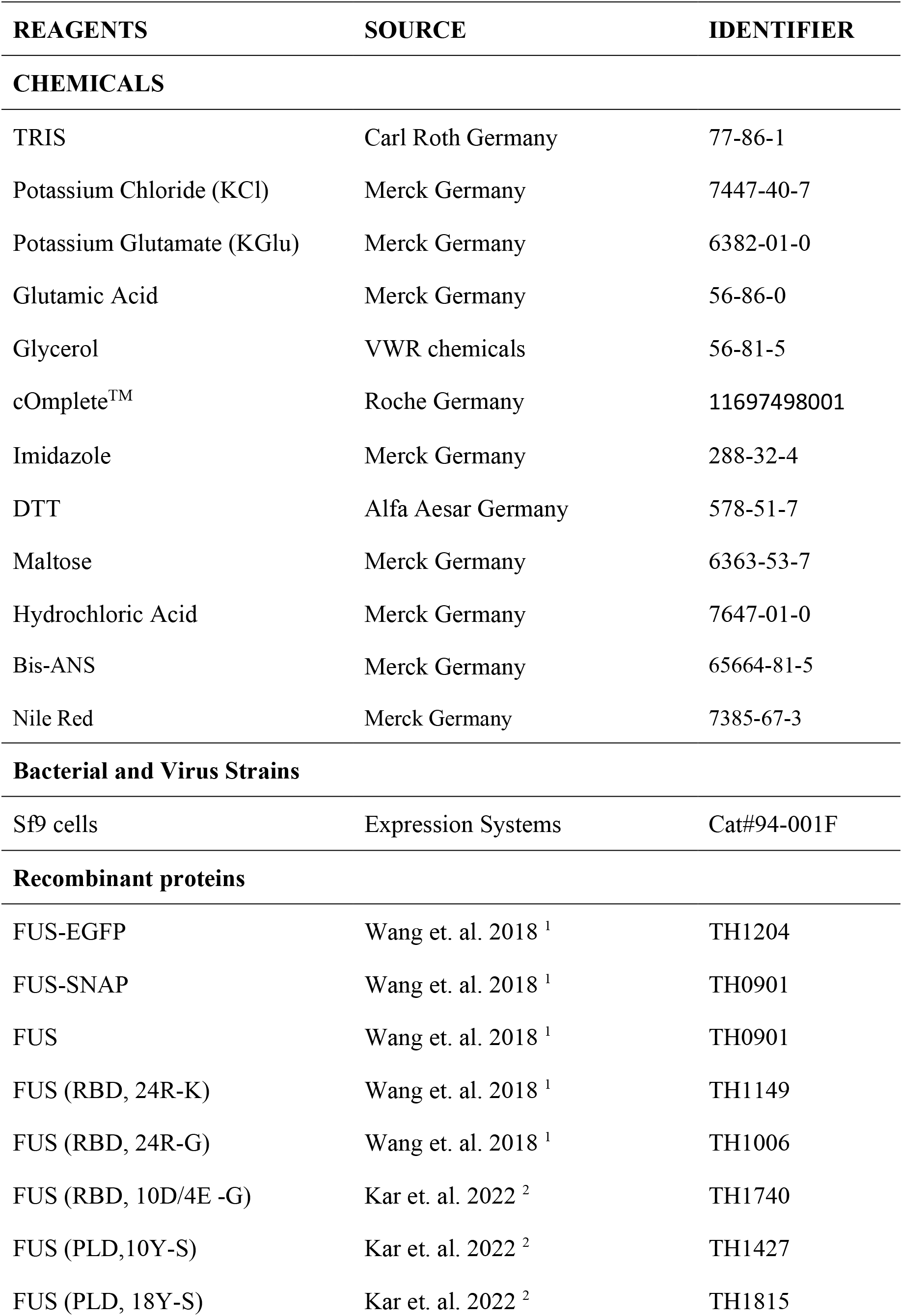
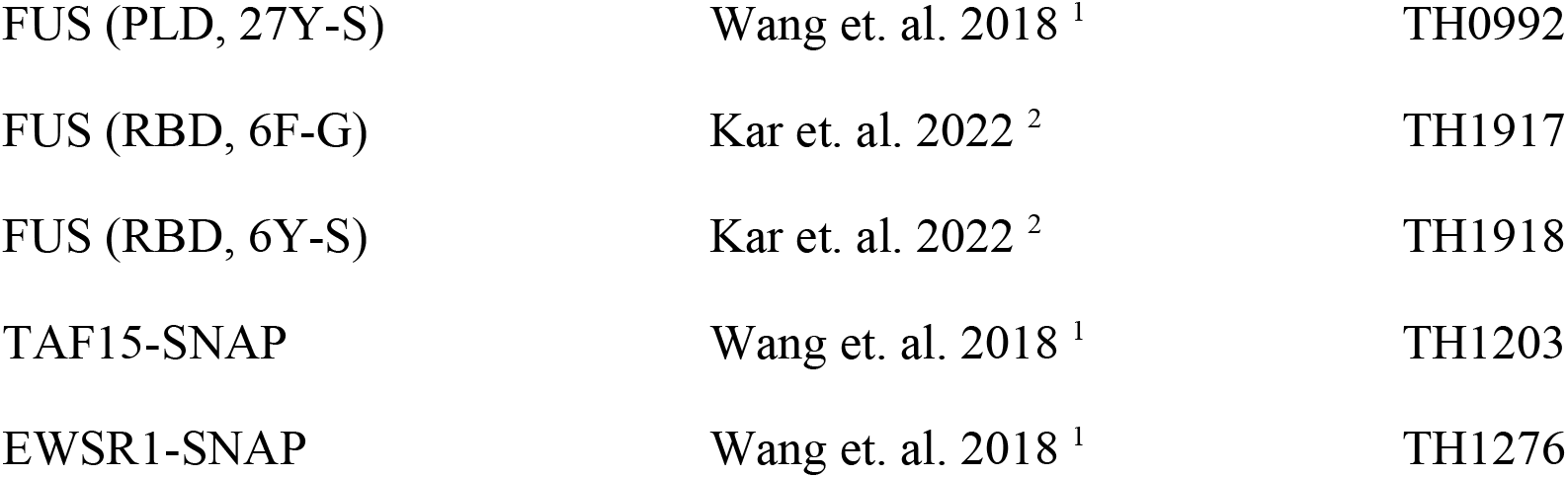
List of reagents, sources, and vendor identifiers if any.

**Table S1:**
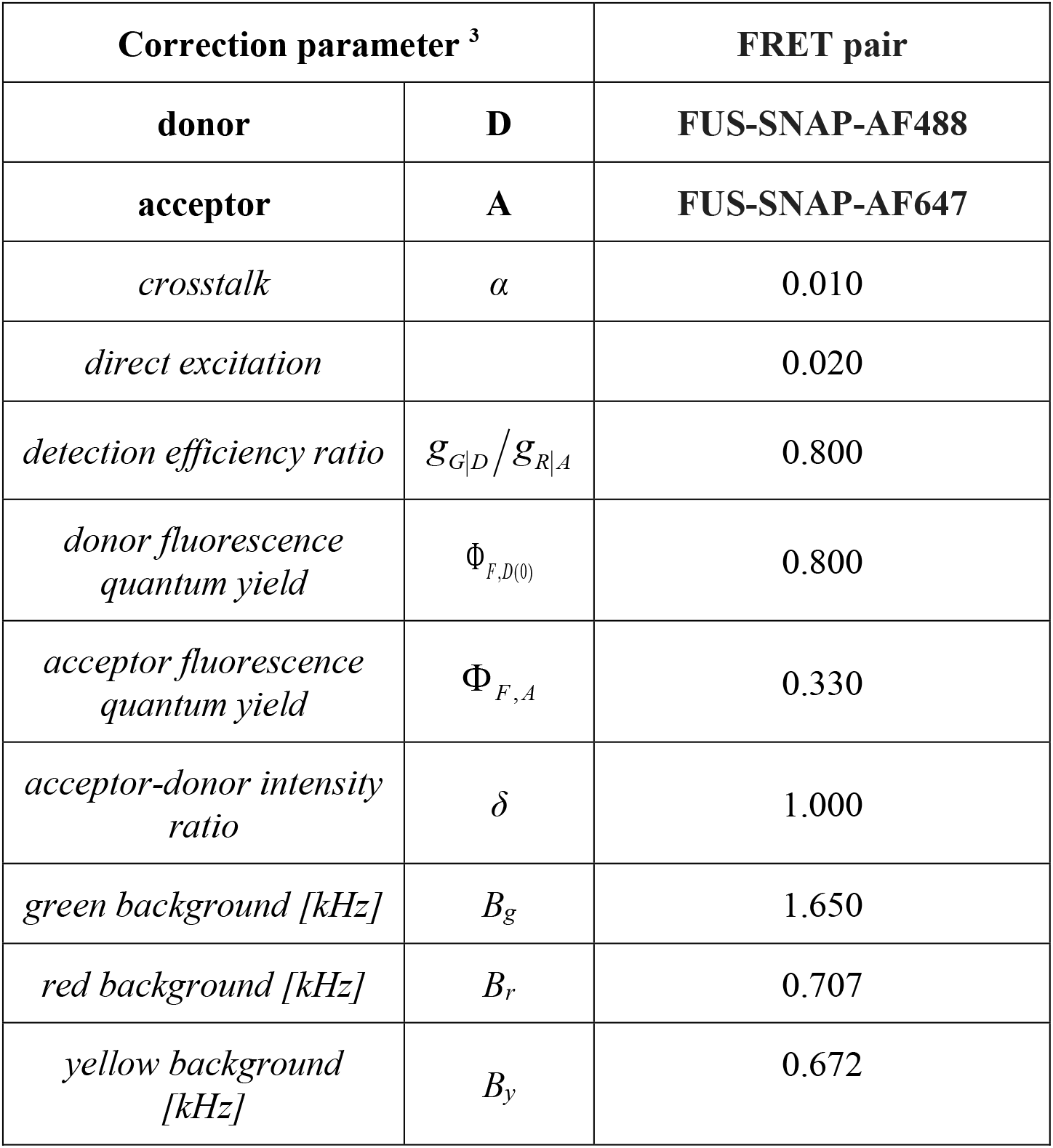
FRET correction parameters.

| Correction parameter <sup>3</sup> |  | FRET pair |
| --- | --- | --- |
| **donor** | **D** | **FUS-SNAP-AF488** |
| **acceptor** | **A** | **FUS-SNAP-AF647** |
| *crosstalk* | $\alpha$ | 0.010 |
| *direct excitation* |  | 0.020 |
| *detection efficiency ratio* | $\mathcal{G}_{G\vert D}/\mathcal{G}_{R\vert A}$ | 0.800 |
| *donor fluorescence quantum yield* | $\Phi_{F,D(0)}$ | 0.800 |
| *acceptor fluorescence quantum yield* | $\Phi_{F,A}$ | 0.330 |
| *acceptor-donor intensity ratio* | $\delta$ | 1.000 |
| *green background [kHz]* | $B_g$ | 1.650 |
| *red background [kHz]* | $B_r$ | 0.707 |
| *yellow background [kHz]* | $B_y$ | 0.672 |

**Table S2:** FCS fit parameters.

| Fit parameter |  | 100 mM KCl | 100 mM KGlu |
| --- | --- | --- | --- |
| $\chi^2$ | *chi-squared* | **1.365** | **40.67** |
| $G_0$ | *correlation offset* | 1.000 | 0.999 |
| $N$ | *number of molecules in focus* | 0.877 | 0.872 |
| $t_{d1,global}$ | *first diffusion time (global) [ms]* | 0.214 | 0.214 |
| $z_{0,1}/\omega_{0,1}$ | *focus ratio for $t_{d1,global}$* | 1.983 | 2.090 |
| $t_{d2}$ | *second diffusion time [ms]* | 1.055 | 2.260 |
| $z_{0,2}/\omega_{0,2}$ | *focus ratio for $t_{d2}$* | 6.125 | 598.4 |
| $R$ | *amplitude for $t_{d1,global}$* | 0.707 | 0.718 |
| $A$ | *amplitude for bunching term* | 0.076 | 0.089 |
| $t_{A,global}$ | *bunching time (global) [ms]* | 0.003 | 0.003 |

### Buffers for the experiments

For buffers, we prepared the following stock solutions;

i. 2 M KCl
ii. 1 M TRIS.HCl pH 7.4 (1 M TRIS was solubilized in DI water and added HCl to adjust the pH to 7.4).
iii. 2 M KGlu
iv. 1 M TRIS.Glu pH 7.4 (1 M TRIS was solubilized in DI water and added Glutamic acid to adjust the pH to 7.4)

### Final buffer composition for protein-based experiments

**100 mM KCl buffer**: The KCl amount was calculated from the residual KCl from the protein stock solution and added to yield a 100 mM final concentration for each experiment. The buffer also contains 20 mM TRIS. HCl pH 7.4.

**KCl-control buffer**: The 1x buffer consists of 20 mM TRIS. HCl pH 7.4 and 100 mM KCl. However, in each experiment, the final KCl was not constant as residual KCl from protein stock was added.

**KGlu buffer**: The 1x buffer consists of 20 mM TRIS. Glu pH 7.4 and 100 mM KGlu. In each experiment, the residual KCl from protein stock was added.

### Constructs, protein expression and purification

The construct/protein sequences used are listed in Section C. Details of materials used for the preparation of samples are as follows:

***Lysis buffer*:** 50 mM TRIS. HCl pH 7.4, 1 M KCl, and 5% Glycerol.

***Protease inhibitor*:** cOmplete™, EDTA-free Protease Inhibitor Cocktail Tablets.

***NTA elution buffer*:** 50 mM TRIS.HCl pH 7.4, 1 M KCl, 5% Glycerol and 300 mM Imidazole.

***MPB elution buffer*:** 50 mM TRIS.HCl pH 7.4, 1 M KCl, 5% Glycerol and 30 mM Maltose.

***Storage buffe*r:** 50 mM TRIS.HCl pH 7.4, 500 mM KCl, 5% Glycerol, and 1 mM DTT.

### Section D. Amino Acid Sequences of Proteins used in Spectroscopic Studies

1. FUS-EGFP: This sequence includes full-length FUS (unshaded), a linker that is cleavable by a TEV protease (shaded in yellow), and the EGFP (shaded in green).

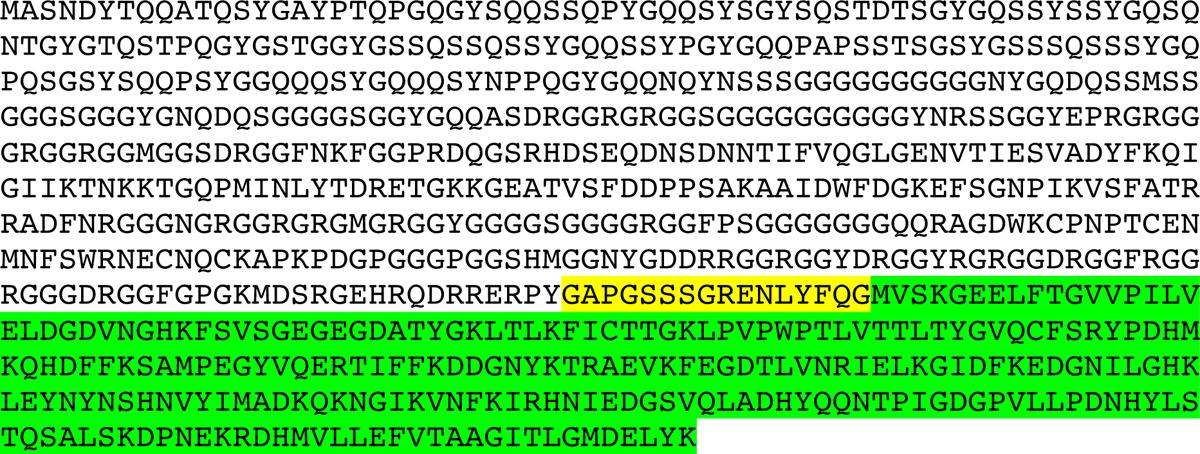

2. FUS-SNAP: This sequence includes full-length FUS (unshaded), a linker that is cleavable by a TEV protease (shaded in yellow), and the SNAP (shaded in gray).

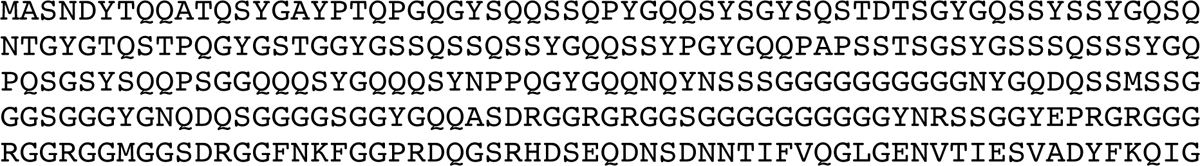
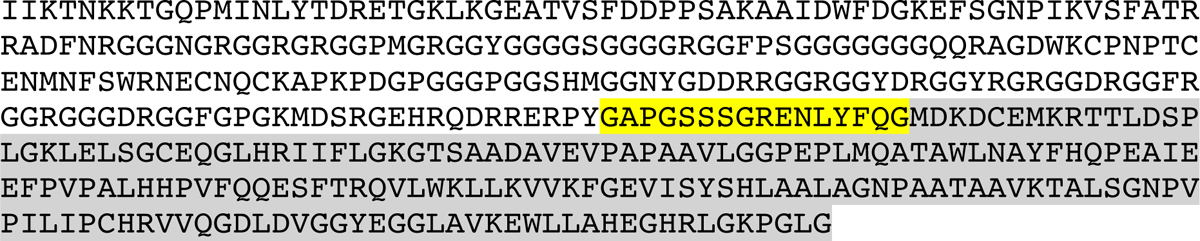

3. FUS

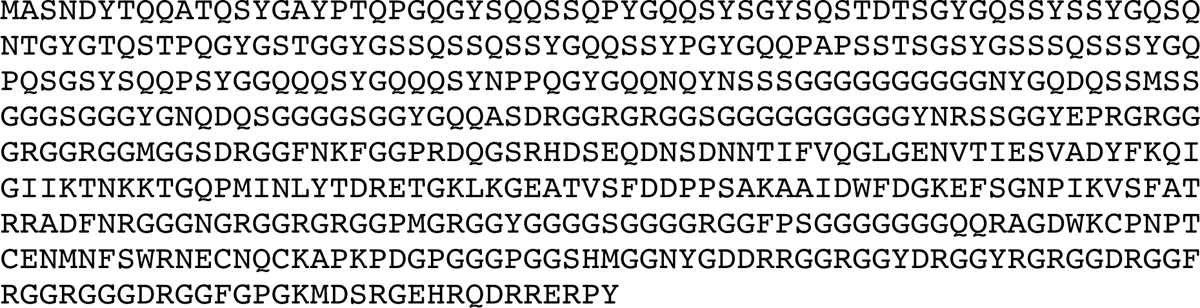

4. FUS(R-K): Full-length FUS with 24 Arg substituted to Lys in the RBD

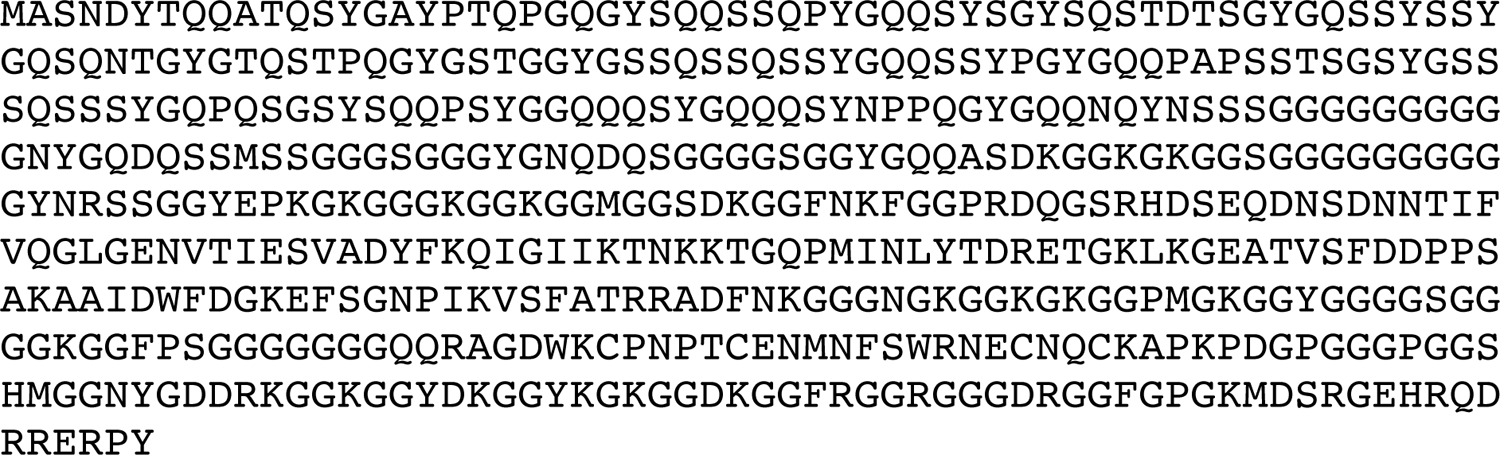

5. FUS(R-G): Full-length FUS with 24 Arg substituted to Gly in the RBD

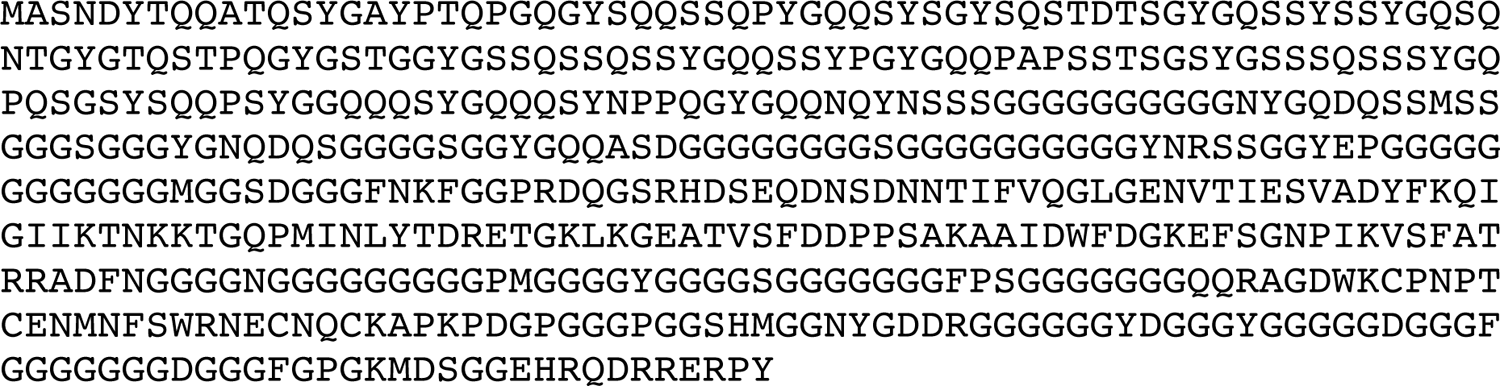

6. FUS-(10D/4E-G): Full-length FUS with 10 Asp and 4 Glu residues substituted to Gly in the RBD

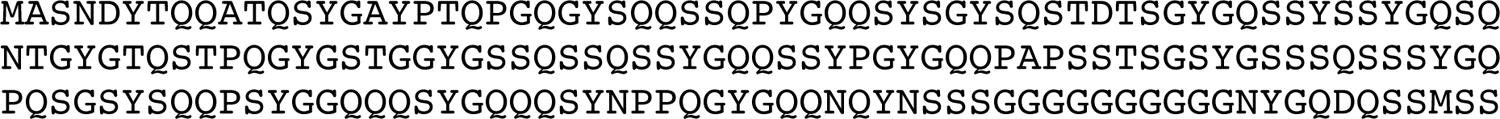
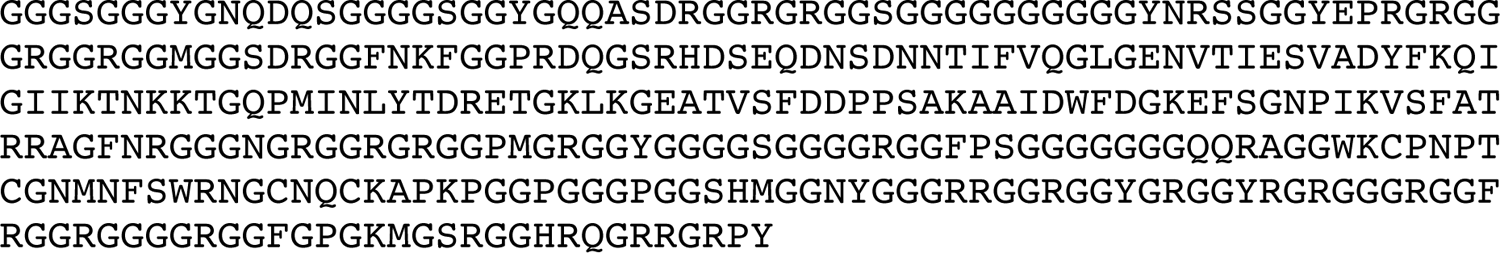

7. FUS(10Y-S): Full-length FUS with 10 Tyr residues substituted to Ser in the PLD

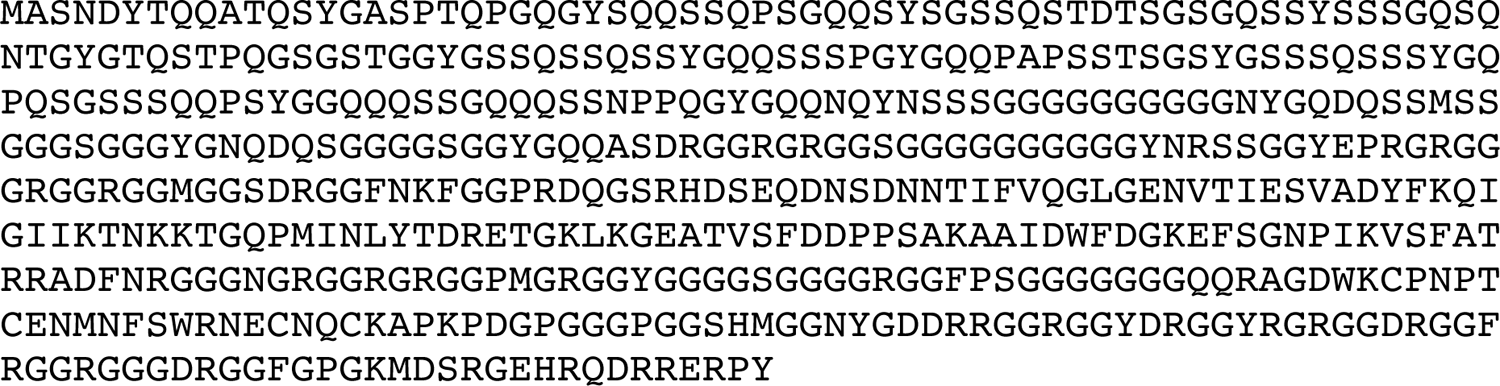

8. FUS(18Y-S): Full-length FUS with 18 Tyr residues substituted to Ser in the PLD

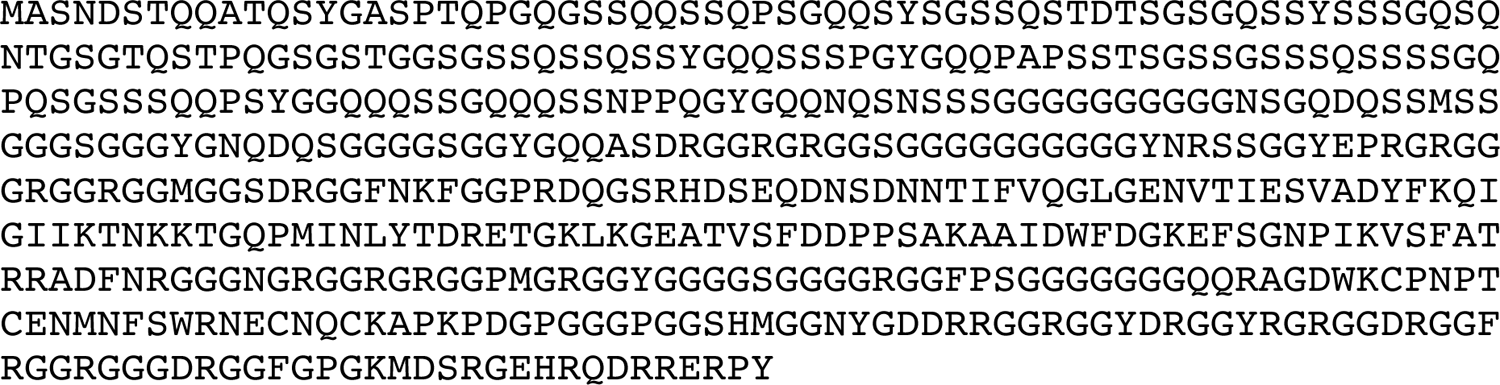

9. FUS(27Y-S): Full-length FUS with 27 Tyr residues substituted to Ser in the PLD

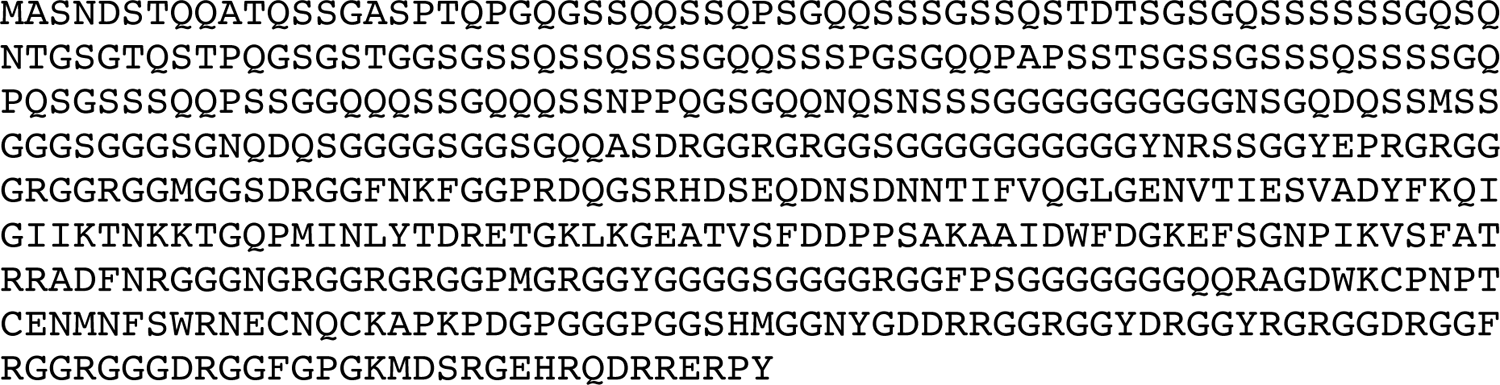

10. FUS(6F-G): Full-length FUS with 6 Phe residues substituted to Gly in the RBD

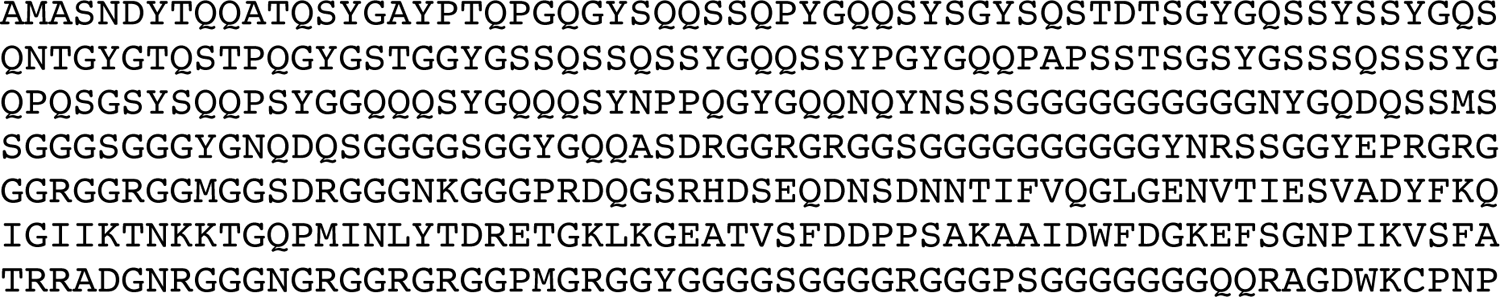
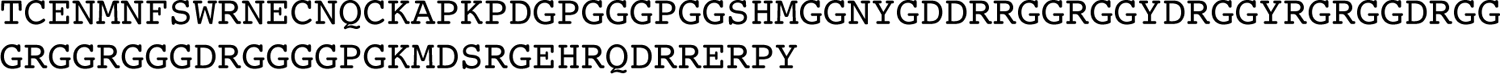

11. FUS(6Y-S): Full-length FUS with 6 Tyr residues substituted to Ser in the RBD

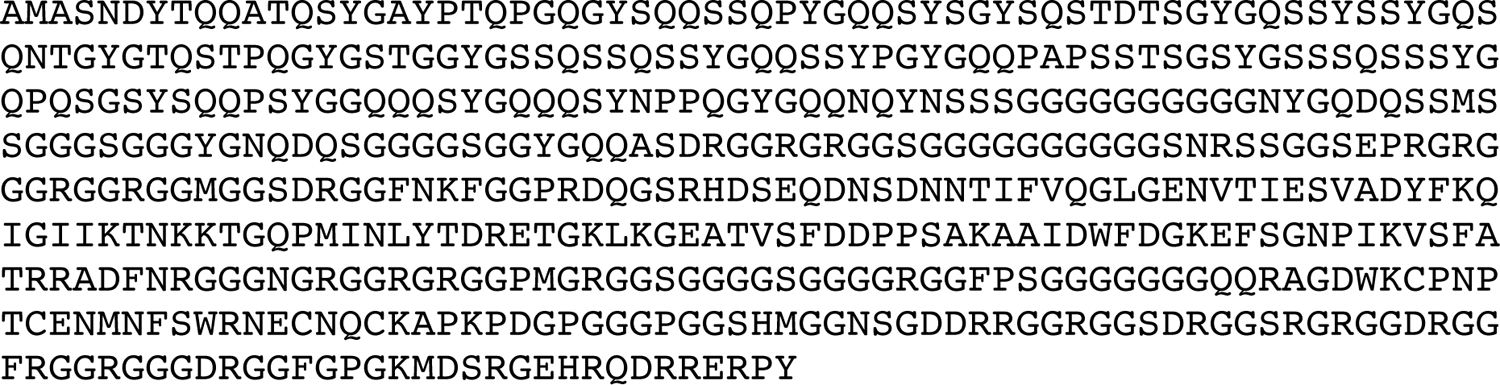

12. Taf15-SNAP: This sequence includes full-length Taf15 (unshaded), a linker that is cleavable by a TEV protease (shaded in yellow), and the SNAP tag (shaded in gray).

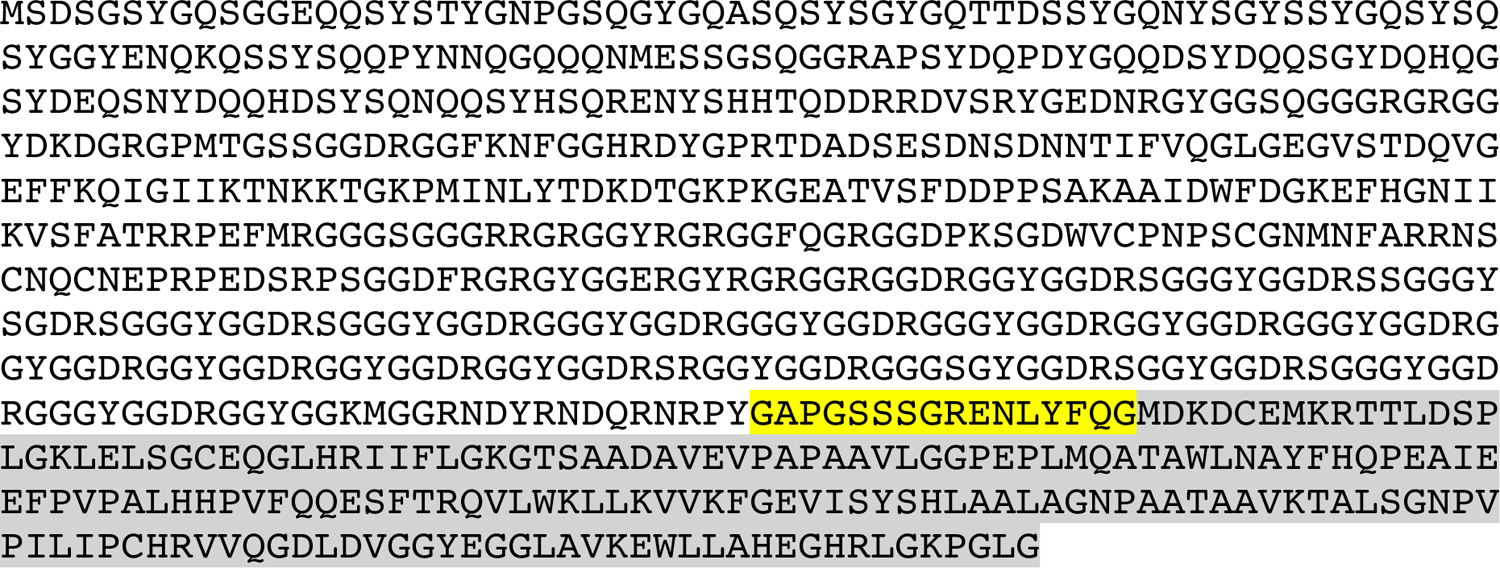

13. Ewsr1-SNAP: This sequence includes full-length Ewsr1 (unshaded), a linker that is cleavable by a TEV protease (shaded in yellow), and the SNAP tag (shaded in gray).

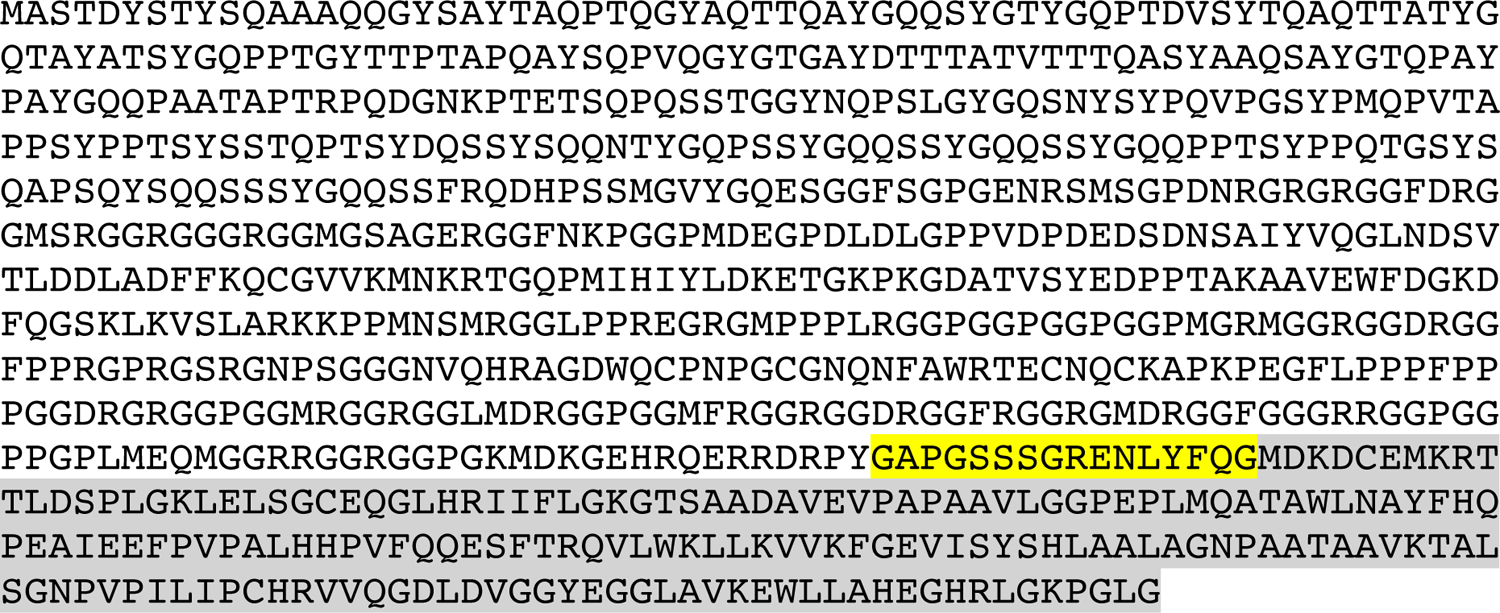

